# Unrestrained ESCRT-III drives chromosome fragmentation and micronuclear catastrophe

**DOI:** 10.1101/517011

**Authors:** Marina Vietri, Sebastian W. Schultz, Aurélie Bellanger, Carl M. Jones, Camilla Raiborg, Ellen Skarpen, Christeen Ramane J. Pedurupillay, Eline Kip, Romy Timmer, Ashish Jain, Philippe Collas, Roland L. Knorr, Sushma N. Grellscheid, Halim Kusumaatmaja, Andreas Brech, Francesca Micci, Harald Stenmark, Coen Campsteijn

## Abstract

The ESCRT-III membrane fission machinery^1,2^ restores nuclear envelope integrity during mitotic exit^3,4^ and interphase^5,6^. Whereas primary nuclei resealing takes minutes, micronuclear envelope ruptures appear irreversible and result in catastrophic collapse associated with chromosome fragmentation and rearrangements (chromothripsis), thought to be a major driving force in cancer development^7-10^. Despite its importance^11-13^, the mechanistic underpinnings of nuclear envelope sealing in primary nuclei and the defects observed in micronuclei remain largely unknown. Here we show that CHMP7, the nucleator of ESCRT-III filaments at the nuclear envelope^3,14^, and the inner nuclear membrane protein LEMD2^15^ act as a compartmentalization sensor detecting the loss of nuclear integrity. In cells with intact nuclear envelope, CHMP7 is actively excluded from the nucleus to preclude its binding to LEMD2. Nuclear influx of CHMP7 results in stable association with LEMD2 at the inner nuclear membrane that licenses local polymerization of ESCRT-III. Tight control of nuclear CHMP7 levels is critical, as induction of nuclear CHMP7 mutants is sufficient to induce unrestrained growth of ESCRT-III foci at the nuclear envelope, causing dramatic membrane deformation, local DNA torsional stress, single-stranded DNA formation and fragmentation of the underlying chromosomes. At micronuclei, membrane rupture is not associated with repair despite timely recruitment of ESCRT-III. Instead, micronuclei inherently lack the capacity to restrict accumulation of CHMP7 and LEMD2. This drives unrestrained ESCRT-III recruitment, membrane deformation and DNA defects that strikingly resemble those at primary nuclei upon induction of nuclear CHMP7 mutants. Preventing ESCRT-III recruitment suppresses membrane deformation and DNA damage, without restoring nucleocytoplasmic compartmentalization. We propose that the ESCRT-III nuclear integrity surveillance machinery is a double-edged sword, as its exquisite sensitivity ensures rapid repair at primary nuclei while causing unrestrained polymerization at micronuclei, with catastrophic consequences for genome stability^16-18^.

---

The formation of dynamic endosomal sorting complex required for transport (ESCRT)-III filaments competent for membrane fission is exquisitely sensitive to balanced levels of the factors that control these filaments^1,2,19^. This is exemplified by depletion of the essential subunit CHMP2A that results in persistent accumulation of CHMP4B, the main component of ESCRT-III filaments, at the reforming nuclear envelope (NE) without successful sealing^3^. We found that these foci are similarly enriched for endogenous CHMP7 and LEMD2 (Fig. 1a, Extended Data Fig. 1a), consistent with the notion that these are key upstream regulators of the ESCRT-III machinery at the NE^3,15^. Considering the delicate balance of ESCRT-III polymerization^19^, we reasoned that overexpression of these targeting factors would phenocopy CHMP2A depletion. Surprisingly, whereas transient or inducible overexpression of CHMP7 rapidly induced CHMP4B foci, these were exclusively cytoplasmic in HeLa and RPE1 cell lines, consistent with reported localization for CHMP7 to the cytoplasm and endoplasmic reticulum (ER)^14,20^ (Fig. 1b top panel, Supplementary Video 1). This suggested that subcellular localization of CHMP7 could be a key determinant to position CHMP4B foci. The CHMP7 C-terminal motif previously annotated as a MIM1 motif^2,21^ closely resembles a type 1a high-affinity nuclear export signal (NES)^22^ (Fig. 1c), suggesting a mechanism to exclude this 45kDa protein from the nucleus^23^. Indeed, fusing this putative CHMP7 NES sequence, but not a mutated version (NES*), to REV-eGFP^24^ was sufficient to relocalize eGFP from nucleoli to the cytoplasm (Fig. 1d, Extended Data Fig. 1b). This was dependent on active export, as assayed by treatment with the Exportin 1 (XPO1) inhibitor leptomycin B (LMB) (Fig. 1d and Extended Data Fig. 1c), indicating that this sequence functions as a *bona fide* NES. Mutating the NES in CHMP7 (CHMP7^NES^*) was sufficient to drive formation of persistent CHMP4B foci associated with the NE as opposed to the ER (Fig. 1b, and Extended Data Fig. 1d-g). Similar results were observed when briefly inducing wild type CHMP7 in the presence of LMB (Fig. 1b) or upon fusion with 2 nuclear localization sequences (CHMP7^NLS^) (Fig. 1b lower panel), arguing against an effect due to functional perturbation of the putative MIM1.

**Figure 1.**
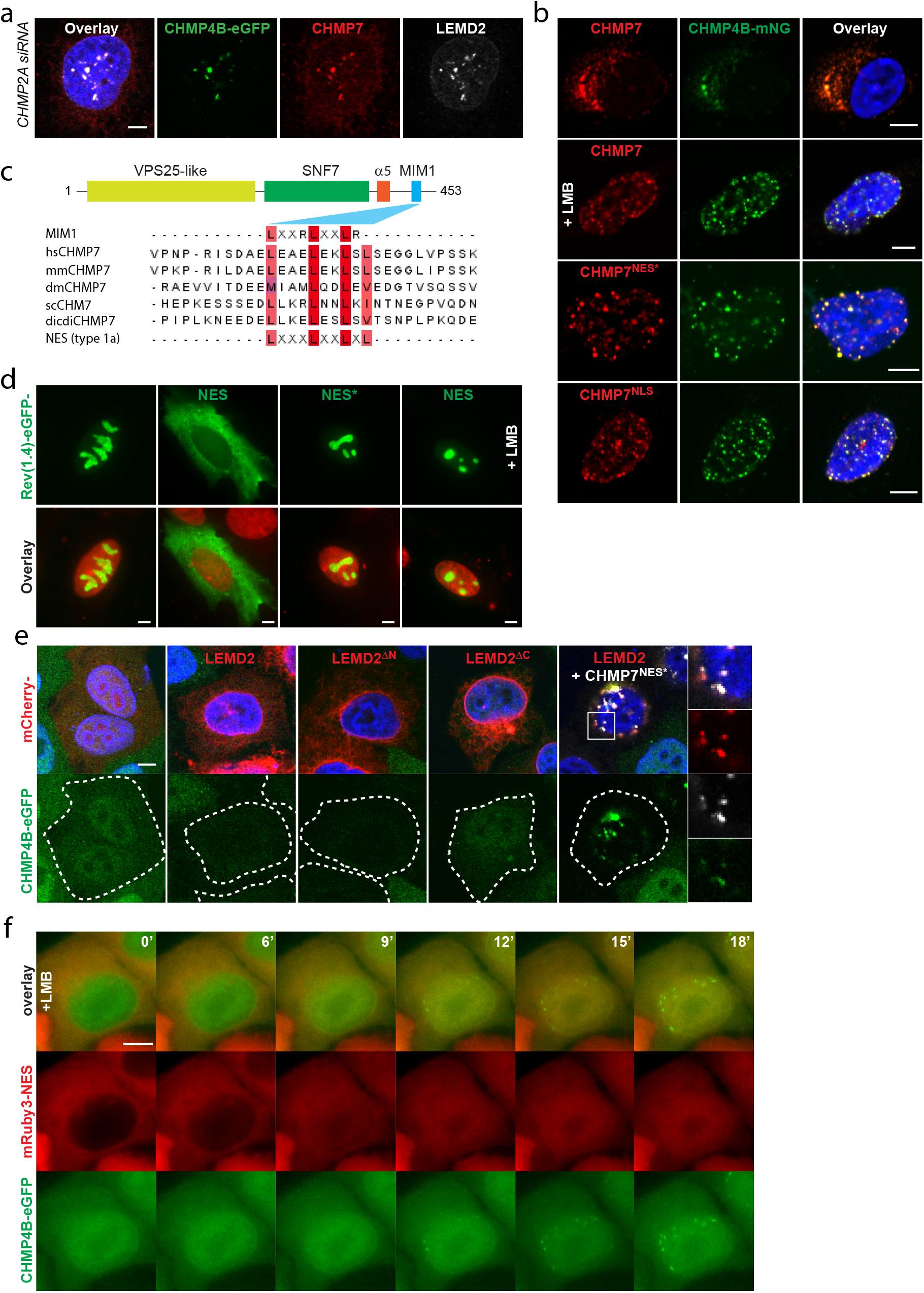
CHMP7 localization governs ESCRT-III activity. **a**, Confocal imaging of endogenous LEMD2 and CHMP7 with CHMP4B-eGFP at the nuclear envelope reveals colocalization. HeLaK CHMP4B-eGFP, LEMD2-SNAP cells were depleted for CHMP2A, fixed and stained as indicated. Scale bar, 5 μm. **b**, Targeting of CHMP4B-mNG foci depend on CHMP7 subcellular localization. RPE1 cell lines stably expressing CHMP4B-mNG were treated with DOX for 2 hours to express the indicated inducible CHMP7-FLAG allele, and with the XPO1 inhibitor leptomycin B (LMB) where indicated, fixed and stained as described. Scale bars, 5 μm. **c**, CHMP7 contains a putative NES. CHMP7 domain structure and sequence alignment of CHMP7 from multiple eukaryotes compared to MIM1 (MIT-interacting motif 1) and type 1a NES consensus sequences. Human (hs), mouse (mm), drosophila (dm), budding yeast (sc), slime mold (dicdi). Conserved residues are shaded. **d**, A CHMP7 c-terminal motif can function as a NES. HeLaK cells were transfected with REV(1.4)-eGFP fusions with aa 421-430 of CHMP7 that constitute a predicted NES, or a mutated version of this sequence (NES*; L421A, L425A, L428A triple mutation), followed by live-cell microscopy. LMB was added where indicated. SiR-Hoechst (red), DNA. Quantification in Extended Data Fig. 1c. Scale bars, 5 μm. **e**, Interplay between LEMD2, CHMP4B and CHMP7 at the nuclear envelope. HeLaK CHMP4B-eGFP cells were transfected with mCherry-fusions of the indicated LEMD2 alleles and with CHMP7^NES^*-FLAG where indicated, fixed, stained and processed for confocal imaging. Outlines are drawn around mCherry positive cells, and the inset is show in the right-hand panel. Scale bar, 5 μm. Quantification of the data in Extended Data Fig. 2f. **f**, Inactivation of XPO1 drives the formation of nuclear CHMP4B foci. HeLaK CHMP4B-eGFP, mRuby3-NES, LEMD2-SNAP cells were treated with LMB and CHMP4B localization was monitored by live-cell imaging, with time after start of imaging indicated. Scale bar, 5 μm. Quantification of the data in Extended Data Fig. 2j.

In contrast, transient overexpression of LEMD2 did not induce CHMP4B foci, but rather resulted in the rapid degradation of nucleoplasmic CHMP4B (Fig. 1e; Extended Data Fig. 2a-c, Supplementary Video 2). This phenotype relied on the MAN1-Src1p C-terminal (MSC) domain of LEMD2 (Fig. 1e, Extended Data Fig.2d,e) that has been reported to mediate interaction with CHMP7^15^. Strikingly, co-expression of LEMD2 with nuclear CHMP7 (CHMP7^NES*^) suppressed CHMP4B degradation and instead resulted in the formation of stable NE foci highly enriched for CHMP4B, CHMP7^NES^*, and LEMD2 (Fig. 1e, and Extended Data Fig. 2f). This phenotype was partially recapitulated in stable LEMD2-SNAPtag^25^, CHMP4B-eGFP cells where CHMP4B reversibly colocalized with LEMD2 enrichments upon nuclear retention of CHMP7 by LMB treatment (Fig. 1f and Extended Data Fig. 2g-j, Supplementary Video 3). These data show that XPO1-mediated export prevents untimely nuclear localization of CHMP7 and argue that nuclear influx of CHMP7 critically controls the formation of stable LEMD2-CHMP7 complexes that license polymerization of the CHMP4B filaments needed for repair^3-6^.

We exploited the subcellular mislocalization of CHMP7^NES^* to monitor effects of excessive nuclear CHMP7 and found that the CHMP4B foci corresponded to areas where the NE underwent architectural deformations, as indicated by enrichment of the ER marker KDEL (Fig. 2a upper panel), the inner nuclear membrane (INM) marker Lap2β, and Lamin A/C (Extended Data Fig. 3a upper and middle panel). The absence of the nucleoporin Nup58 (Fig. 2a lower panel) argued against these foci representing aggregates of aberrant nuclear pore complexes^26,27^. Correlative light and electron microscope (CLEM) analysis revealed these membrane distortions to be complex 3-dimensional networks with trabecular appearance (Fig. 2b). Whereas expression of CHMP7 resulted in remodelling of the ER without much effect on the NE (Fig.2b, left panel), CHMP7^NLS^ selectively distorted the INM (Fig. 2b, right panel), with CHMP7^NES^* evolving a composite phenotype affecting both ER and INM (Fig. 2b, middle panel). Additionally, LMB treatment induced nuclear CHMP4B foci that transformed LEMD2-enriched parallel INM sheets into 3-dimensional membrane networks (Extended Data Fig. 3b). Considering the dramatic membrane deformation and the absence of Lamin B1 at these sites (Extended Data Fig. 3a lower panel)^6,7,28^, we assessed NE integrity upon expression of these CHMP7 alleles by monitoring nuclear exclusion of an mRuby3-NES compartmentalization marker. Whereas overexpression of LEMD2 or CHMP7 did not compromise nuclear integrity, CHMP7^NLS^ expressing cells lost compartmentalization within a few hours (Fig. 2c). Cells expressing CHMP7^NES^* also decompartmentalized albeit slower than CHMP7^NLS^, in a fashion dependent on its membrane-binding VPS25-like domain^14^ (Fig. 2c and Extended Data Fig. 3 c, d). This was not due to defective mitotic NE reformation as it was readily observed in cells that did not traverse mitosis (Supplementary Video 1). Taken together, these data show that persistent ESCRT-III foci formed upon excessive nuclear influx of CHMP7 deform the NE to the point of irreversible rupture.

**Figure 2.**
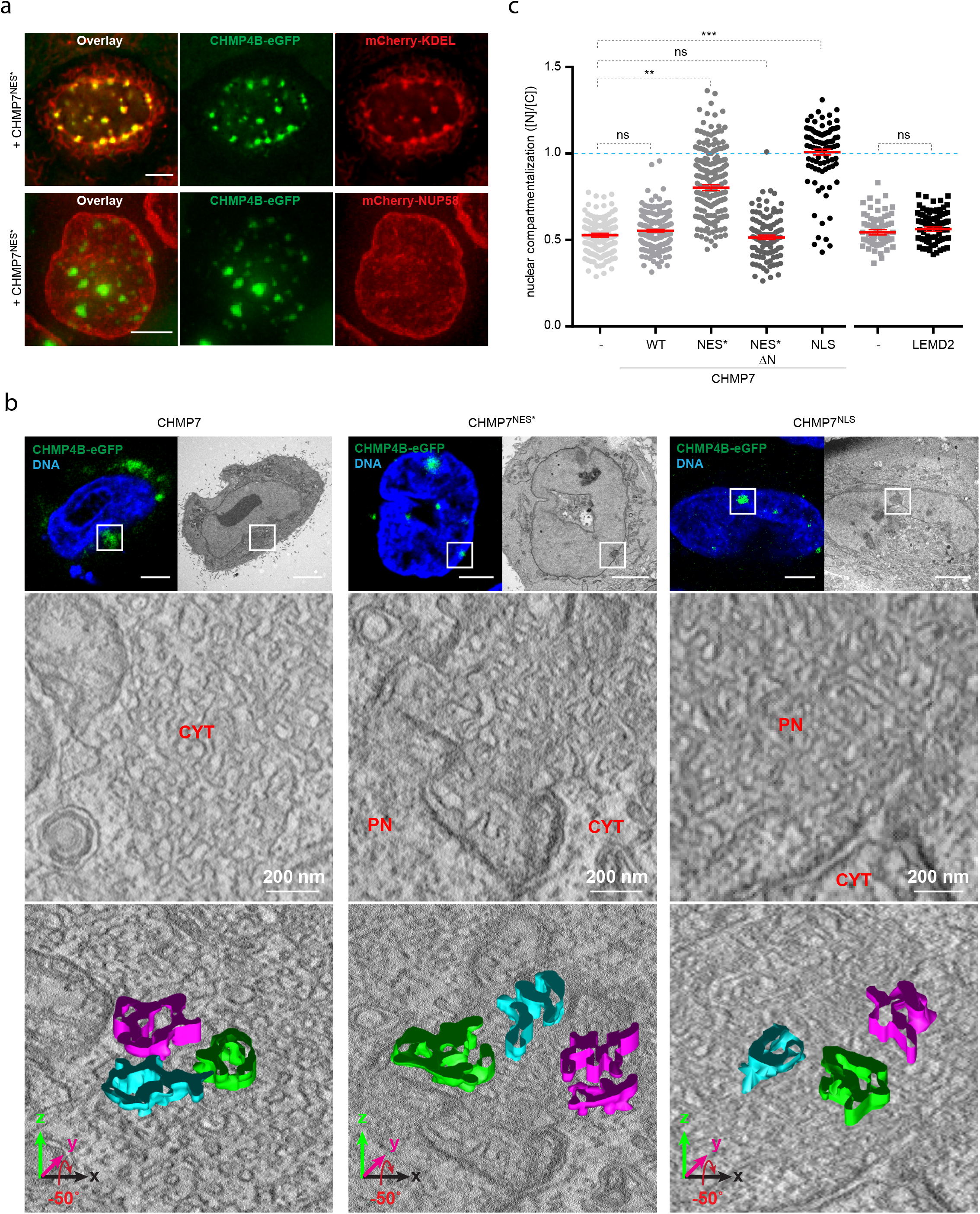
Unrestrained nuclear CHMP7 drives NE deformation and rupture. **a**, Persistent CHMP4B foci colocalize with membrane clusters but not with nuclear pore complexes. HeLaK CHMP4B-eGFP cells stably expressing mCherry-KDEL or NUP58-mCherry were transiently transfected with CHMP7^NES^*-FLAG and imaged by live-cell imaging. Scale bars, 5 μm. **b**, CLEM tomography analysis shows that formation of persistent CMP4B foci results in extensive membrane deformation. HeLaK CHMP4B-eGFP, mRuby3-NES cells were transiently transfected with indicated CHMP7 alleles, followed by imaging, fixation and processing for electron microscopy (top and middle panels). Bottom panels, the tilted representation with modelling of 3 isolated sections of membrane. PN, primary nucleus; Cyt, cytoplasm. Scale bars, 5 μm. **c**, Overexpression of nuclear CHMP7 results in irreversible membrane rupture. HeLaK CHMP4B-eGFP, mRuby3-NES cells were transfected with indicated constructs and nucleo-cytoplasmic compartmentalization of transfected cells was measured after 24h. Bars, mean and SEM from 3 experiments. n= 217, 172, 180, 102, 88, 62, 82. **P=0.0067; ***P=0.0004 twotailed unpaired *t*-test, df=4.

Whereas LEMD2 overexpression alone depleted all nuclear CHMP4B, CHMP4B associated with micronuclei (MN) was refractory to this phenomenon (Fig. 3a). Instead, these MN were highly enriched for endogenous CHMP7, suggestive of nuclear accumulation of CHMP7 (Fig. 3a). Similarly, a fraction (2.48% ± 0.36% SEM) of MN showed striking enrichment of CHMP4B-eGFP, endogenous LEMD2 and CHMP7 (Fig. 3b and Extended Data Fig. 4 a,b), as well as other ESCRT-III subunits (Extended Data Fig. 4c). Reasoning that these could represent MN with compromised MN envelope (MNE) integrity, we monitored CHMP4B dynamics at monopolar spindle 1 (MPS1)-inhibitor (AZ3146) induced MN as they underwent rupture. Following rupture, CHMP4B accumulated at the MNE within minutes, and this recruitment was reduced for a membrane-binding defective CHMP4B^4DE^ mutant^29^ (Fig. 3c, Extended Data Fig. 4d-f, Supplementary Videos 4,5). Together with similar CHMP4B recruitment kinetics to primary nuclei (PN) and MN upon CHMP7^NES^* expression (Extended Data Fig. 4g), this argues that inherent defects in MN^30^ do not extend to ESCRT-III targeting. However, rather than the characteristic transient recruitment associated with successful repair^3,5^, CHMP4B progressively accumulated in multiple coalescing foci over hours, without any detectable MNE repair (Fig. 3c, Supplementary Video 4,5). This was associated with a dramatic increase of LEMD2 levels spreading from the site of rupture (Fig. 3d, Supplementary Video 6), a phenotype that depended on CHMP7 but less on CHMP4B (Fig. 3e, Supplementary Video 7). Subsequently, the MNE rapidly changed morphology from being smoothly round to becoming progressively more convoluted and condensed (Fig. 3f, Supplementary Video 7,8), most notably at sites of CHMP4B accumulation (Extended Data Fig. 4h). Depletion of CHMP4B or CHMP7 completely reverted this morphology in ruptured MN (Fig. 3f and Extended Data Fig. 4i). CLEM experiments consistently revealed ESCRT-III dependent complex trabecular membrane networks at sites of CHMP4B accumulations in ruptured MN (Fig. 3g, Extended Data Fig. 5, Supplementary Video 9). These complex 3-dimensional membrane networks showed striking architectural similarities to those observed at PN upon forced mislocalization of CHMP7 (Fig. 2b, Extended Data Fig. 6). This demonstrates a causal role for unrestrained ESCRT-III accumulation in driving MN catastrophe following MN rupture. As depletion experiments showed that ESCRT-III did not drive MN rupture (Extended Data Fig. 4j,k), it raises the interesting possibility that it generates secondary ruptures at sites of membrane distortion as the MN collapses (Fig. 2a-c and Extended Data Fig. 4h).

**Figure 3.**
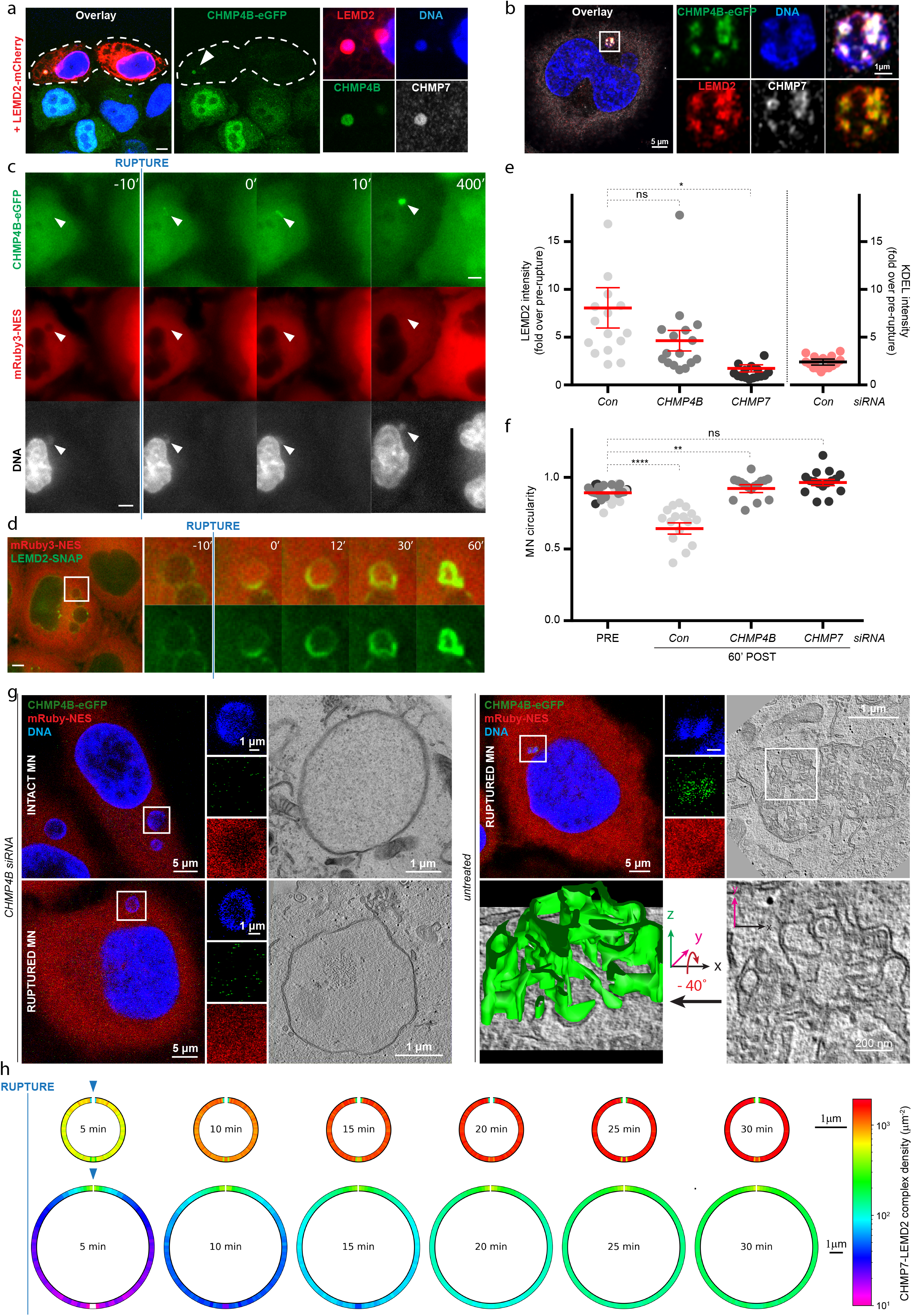
Ruptured MN lack the ability to restrain ESCRT-III activity and undergo ESCRT-III driven catastrophe. **a**, Micronuclear CHMP4B is resistant to LEMD2 overexpression induced depletion. HeLaK CHMP4B-eGPF cells were transfected with LEMD2-mCherry, fixed, stained for endogenous CHMP7 and imaged by confocal microscopy. LEMD2-mCherry overexpressing cells are outlined, and a MN is indicated (arrowhead). Scale bar, 5 μm. **b**, CHMP4B, LEMD2 and CHMP7 enrich at a fraction of MN. HeLaK CHMP4B-eGFP cells were treated with AZ3146 fixed, stained as indicated and imaged by Airyscan microscopy. **c**, CHMP4B persistently accumulated at MN upon rupture. HeLaK CHMP4B-eGFP, mRuby3-NES cells were treated with AZ3146 to induce formation of MN, stained with SiR-Hoechst, and events at MN were monitored by live cell imaging. Arrowheads indicate a rupturing MN. Scale bar, 5 μm. **d**, LEMD2 hyperaccumulates at MN upon rupture. HeLaK mRuby3-NES, LEMD2-SNAP cells were treated with AZ3146 to induce formation of MN, stained with SiR-SNAP, and events at MN were monitored by live cell imaging. Scale bar, 5 μm. **e**, LEMD2 hyperaccumulation depends on CHMP7 and CHMP4B. As with panel d, but including treatment with indicated siRNAs. LEMD2 intensity at the micronuclear envelope (MNE) was monitored by live-cell imaging, intensity fold increase 60 minutes after rupture was quantified and plotted. Intensity changes in mCherry-KDEL are included as reference. Bars indicate mean and SEM of 3 experiments for LEMD2, mean and 95% confidence interval for KDEL. Con n=15; CHMP4B n=16; CHMP7 n=15; Con KDEL n=17. ***P=0.0001 two-tailed unpaired *t*-test with Welch’s correction. **f**, CHMP4B accumulation results in collapse of the MNE. HeLaK CHMP4B-eGFP, mCherry-KDEL cells were treated with indicated siRNAs, and MN were monitored by live-cell imaging for rupture events. MN circularity was measured before rupture and 60 minute after rupture, with circularity a proxy for membrane deformation. Bars indicate mean and SEM from 3 experiments. n=16 each condition. ***P <0.0001, **P=0.0024 paired *t*-test, df= 4. **g**, CHMP4B hyperaccumulation at ruptured MN induces membrane deformations similar to those observed at PN. HeLaK CHMP4B-eGFP, mRuby3-NES cells were treated with indicated siRNAs, incubated with AZ3146 to induce MN, fixed and imaged by confocal microscopy to identify ruptured MN. Samples were processed for correlative electron microscopy. A 3D-model of the MNE was built from electron tomography of a untreated ruptured MN (left panel). **h**, CHMP7-LEMD2 complex accumulation along the INM of MN (top panels) and PN (bottom panels) following rupture. Simulation experiments describing interaction dynamics of cytoplasmic CHMP7 and INM-bound LEMD2 in PN (400 μm^3^) and MN (4 μm^3^) at different time points following NE rupture (100nm, at the north pole, indicated by arrowhead) while considering influx of novel LEMD2 molecules (maximum occupancy fraction 0.5; Extended Data Fig. 7a). A five-color heatmap was used to visualize the local density of CHMP7-LEMD2 complexes along the INM, as a function of the angle from the pore (located at the top of the figure). The parameters used for simulations are described in the Materials and Methods section. Scale bars, 1 μm.

CHMP7-dependent spreading of LEMD2 accumulation across the MNE (Fig. 3e) suggested that MN lack the capacity to restrict CHMP7-LEMD2 complexes to the site of rupture, thus licensing unrestrained formation of ESCRT-III filaments at sites along the MNE. To test this hypothesis, we performed in silico experiments of NE rupture at PN and MN using INM concentrations of LEMD2-CHMP7 complexes as readout (Extended Data Fig. 7, materials and methods). Upon rupture, continuous influx of cytoplasmic CHMP7 rapidly overloaded the MN pool of LEMD2 along the entire MNE (Fig. 3h top panel), while at PN CHMP7-LEMD2 accumulation remained restricted to the site of rupture. Including influx of new LEMD2 molecules upon saturation (Extended Data Fig. 7a), the concentration of LEMD2-CHMP7 complexes at MN far exceeded that at PN over the course of 30 minutes, a time range reflecting physiological nuclear envelope repair (Fig. 3h). Similar differences between MN and PN were observed when assuming CHMP7 exclusively membrane-associated as opposed to cytoplasmic^14,20^ (Extended Data Fig. 7b,c). Considering the delicate balance governing ESCRT-III function^19^, these simulations suggest that MN inherently lack the capacity to restrict CHMP7-LEMD2 complexes to sites of rupture and to the physiological concentration required to drive membrane repair. Excessive CHMP7-LEMD2 complexes subsequently license unrestrained CHMP4B polymerization at numerous sites along the MNE, diverting ESCRT-III machinery activity from rapid membrane fission to persistent membrane distortion.

Considering the ESCRT-III induced morphological changes at PN and ruptured MN, we explored the relation between ESCRT-III accumulation and the stability of the underlying genome. We exploited ectopic nuclear localization of CHMP7^NES^* to PN as a tractable model. CHMP7^NES^*-induced nuclear ESCRT-III foci were associated with rosettes of γH2Ax (Extended Data Fig. 8a), and were enriched for the single-stranded DNA (ssDNA)-binding replication protein A2 (RPA2)^31^ (Fig. 4a). The facts that RPA2 enrichment was restricted to CHMP4B foci at the nuclear surface (Manders’ colocalization coefficient RPA/CHMP4B 0.79 ± 0.23 SD) and that RPA2 was apparent even in the absence of NE ruptures (Extended Data Fig. 8b,c) argued against influx of cytoplasmic stressors underlying these phenotypes. Because of the dramatic membrane distortions combined with accumulation of chromatin-associated proteins like LEMD2^32^, we investigated whether RPA2 foci corresponded to areas subject to DNA torsional stress^33^. Strikingly, we found that Topoisomerase IIb (Top2B), the enzyme that relieves torsional stress in chromatin topological domains by transiently generating chromosome breaks^34^, was highly enriched at these foci (Fig. 4b and Extended Data Fig.8d). As Top2B has previously been linked to chromosome rearrangements^34^ we tested the consequences of this torsional stress on chromosome integrity by performing metaphase spreads following induction of CHMP7 alleles. Whereas overexpression of wild-type CHMP7 induced a low-level of DNA breaks, induction of CHMP7^NES*^ dramatically increased the frequency of chromosome breaks and fragmentation (Fig. 4c,d).

**Figure 4.**
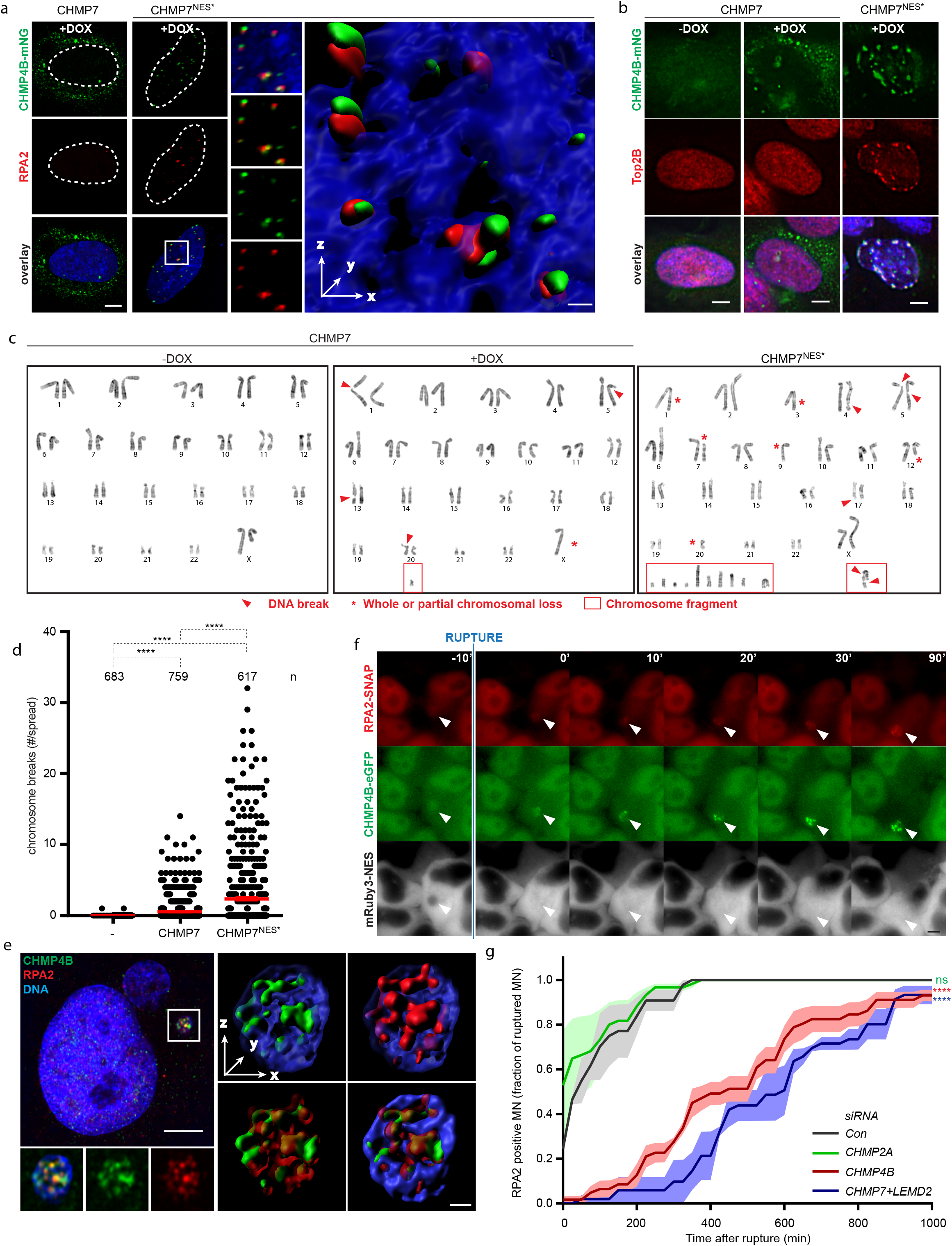
Unrestrained CHMP7 drives DNA damage, torsional stress and chromosome fragmentation. **a**, Nuclear CHMP4B foci associate with the ssDNA marker RPA2. RPE1 CHMP4B-mNG, mRuby3-NES and inducible CHMP7-FLAG or CHMP7^NES^*-FLAG cells were treated with DOX to induce the indicated CHMP7 allele, fixed, stained for RPA2 and imaged by Airyscan microscopy. Nuclei are outlined. An Imaris surface 3D-rendering (right panel) of RPA2 foci localizing at or in close proximity to CHMP4B foci at the surface of the PN (inset). For n=12 PN, Manders’ colocalization coefficient RPA/CHMP4B 0.79 ± 0.23. Scale bars, 5 μm and 0.3 μm (Imaris reconstruction). **b**, Nuclear CHMP7 induces DNA torsional stress. RPE1 CHMP4B-mNG, mRuby3-NES and inducible CHMP7-FLAG or CHMP7^NES^*-FLAG cells were treated with DOX to induce the indicated CHMP7 allele, fixed, stained for Top2B and imaged by confocal microscopy. Quantification of Top2B accumulation in Extended Data Fig. 8d. Scale bars, 5 μm. **c.** Nuclear CHMP7 drives chromosome breaks and fragmentation. RPE1 mRuby3-NES and inducible CHMP7-FLAG or CHMP7^NES^*-FLAG cells were treated with DOX to induce the indicated CHMP7 allele, incubated overnight with colchicine and processed for cell harvesting and metaphase spread analysis. Representative examples are shown. **d**, Quantification of the experiments described in panel c, with n indicated for each condition. Bars, mean and SEM. ****P < 0.0001 two-tailed Mann-Whitney test. Control (-) n=683; CHMP7 n=759; CHMP7^NES^* n=617. **e**, CHMP4B colocalizes with RPA2 at ruptured MN. HeLaK cells were treated with AZ3146 to induce MN, fixed, stained as indicated and imaged by Airyscan microscopy. CHMP4B and RPA2 show a strong spatial correlation. For n=14 MN, Pearson’s correlation 0.84 ± 0.15; Manders’ colocalization coefficient RPA/CHMP4B 0.94 ± 0.05. Right panel, Imaris surface 3D-rendering. Scale bars, 5 μm and 0.5 μm (Imaris reconstruction). **f**, RPA2 enrichment at ruptured MN follows CHMP4B foci formation. HeLaK CHMP4B-eGFP, mRuby3-NES, RPA2-SNAP cells were incubated with AZ3146, stained with SiR-SNAP, and resulting MN were monitored by live-cell imaging. Accumulation of RPA2 is indicated relative to time of rupture. Arrowhead indicates a rupturing MN. Scale bar, 5 μm. **g**, RPA accumulation at ruptured MN requires ESCRT-III. HeLaK CHMP4B-eGFP, mRuby3-NES, RPA2-SNAP cells were treated with indicated siRNAs, incubated with AZ3146, stained with SiR-SNAP and resulting MN were monitored every 5 min by live-cell imaging. MN are considered RPA2-positive when RPA enrichment is observed following rupture, and fraction of RPA2-positive ruptured MN is plotted over time. Mean (line) and SEM (bands) are indicated. Con n=41; CHMP2A n=46; CHMP4B n=43; CHMP7+LEMD2 n=27. ****P< 0.0001 two-tailed unpaired *t*-test. Df=8 (CHMP4B vs Con); Df=7 (CHMP7+LEMD2 vs Con).

Reasoning that this same phenomenon could drive chromosome fragmentation in ruptured MN, we monitored RPA2 accumulation at MN. Indeed, RPA2 colocalized with CHMP4B foci at MN (Manders’ colocalization coefficient RPA/CHMP4B 0.94 ± 0.05 SD). Live-cell imaging showed that upon MN rupture RPA2 accumulated rapidly at sites of CHMP4B enrichment along the MNE (Fig. 4e,f and Extended Data Fig. 8e, Supplementary Video 10). Micronuclear RPA2 accumulation required unrestrained ESCRT-III driven membrane distortion as depletion of CHMP4B or CHMP7 and LEMD2 suppressed RPA2 accumulation without restoring MN integrity (Fig. 4g, Supplementary Video 10). These data show that unrestrained accumulation of ESCRT-III at PN and MN has catastrophic effects on the underlying DNA, inducing torsional stress, the generation of ssDNA and chromosome fragmentation.

Taken together our data suggest a compartmentalization sensing mechanism for loss of NE integrity. Rupture of the NE permits local nuclear influx of CHMP7 and its association with LEMD2 to license ESCRT-III activation required for sealing (Extended Data Fig. 9). The inherent weakness of this system is highlighted at MN, which lack the capacity to restrict CHMP7-LEMD2 complexes to the site of rupture, and instead allow unrestrained activation of ESCRT-III along the surface of the INM (Extended Data Fig. 9). This phenomenon is associated with dramatic membrane distortion and consequent DNA torsional stress, formation of ssDNA, chromosome damage, as well as the recruitment of exonucleases such as TREX1 (Extended Data Fig. 8f-h) ^35,36^. This ultimately drives the fragmentation of chromosomes, possibly in combination with stress ensuing from defective DNA replication and premature chromosome condensation frequently associated with ruptured micronuclei^7,9^. In conclusion, our work highlights the ESCRT-III machinery as a double-edged sword, and suggests that it presents a conditional non-genetic driver of genome instability and the development of cancer.

## Author contributions

M.V. and C.C. conceived and designed the study with important contributions from H.S. and S.W.S.. M.V. generated cell lines, performed light microscopy, analysed data, and prepared figures. S.W.S. performed and analysed EM experiments, and prepared figures. A.Be. generated cell lines, performed light microscopy, analysed data, and prepared figures. C.M.J. performed and analysed the simulation experiments, and prepared figures. C.R., E.S., E.K., R.T., and A.J. performed light microscopy and analysed data. C.R.J.P. performed metaphase spreads experiments. P.C. provided comments and infrastructure. R.L.K. provided comments and contributed to the simulation setup. S.N.G. and H.K. supervised the simulation experiments. A.Br. supervised and performed EM experiments and provided conceptual input. F.M. supervised and analysed the metaphase spreads experiments. H.S. provided infrastructure and co-supervised the study. C.C. supervised the study, generated constructs and cell lines, performed light microscopy, analysed data, and prepared figures. M.V. and C.C. wrote the paper with contributions from all authors.

## Acknowledgements

We thank U. Dahl Brinch, M. Smestad and L. Kymre for assistance with EM, K. Schink for cell lines, E. Rønning, A. Bergersen, and A. Baumeister for technical support, A. Engen for assistance with cell culture, and C. Bassols for IT support. We also thank V. Nähse, T. Stokke, R. Syljuåsen, T. Liyakat Ali for valuable discussions. The National Core Facility for Human Pluripotent Stem Cells, the Advanced Light Microscopy Core Facility (Oslo University Hospital), the Advanced Electron Microscopy Core Facility (Oslo University Hospital), the Norwegian Advanced Light Microscopy Imaging Network (University of Oslo), and the Section for Cancer Cytogenetics (Oslo University Hospital) are acknowledged for providing facilities. C.C. was supported by the Research Council of Norway (project number 262375). H. S. was supported by the Norwegian Cancer Society (project number 182698) and the South-Eastern Norway Regional Health Authority (project number 2016087). This work was partly supported by the Research Council of Norway through its Centres of Excellence funding scheme (project number 262652).

## Legends to Extended Data Figures

**Extended Data Figure 1.**
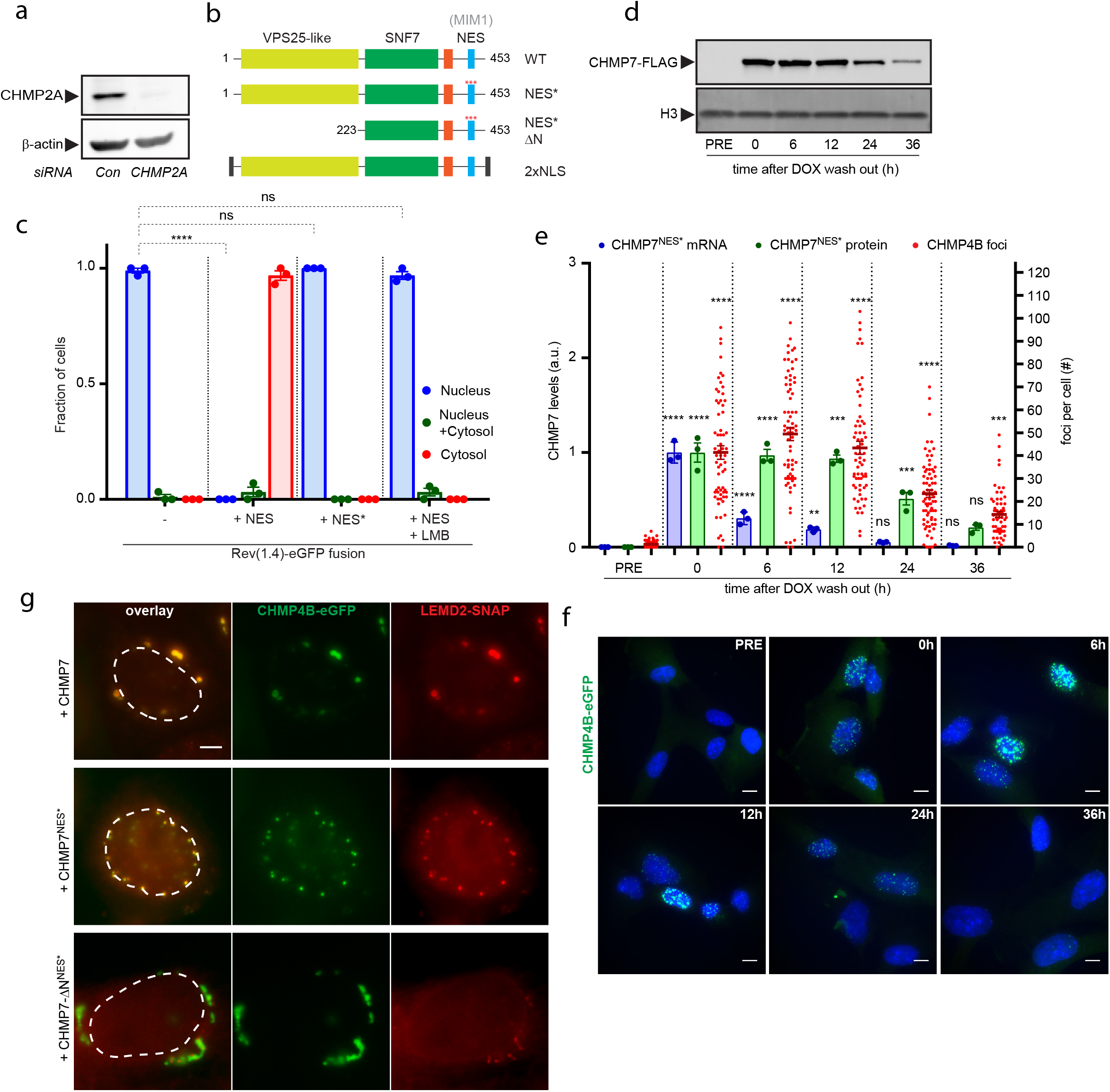
CHMP7 is excluded from the nucleus through a NES. **a**, Immunoblot of whole-cell lysate of HeLaK CHMP4B-eGFP cells showing efficient depletion of CHMP2A upon siRNA treatment. **b**, Schematic representation of human CHMP7 functional domains and mutant alleles used in this study. **c**, Quantification of subcellular localization of REV(1.4)-eGFP fusions as shown in Fig. 1d. LMB treatment where indicated. Bars, SEM, dots represent the mean of each independent experiment. n= 218, 149, 138, 153 cells. ****P<0.0001, two-tailed unpaired t-test, df=4. **d**, RPE1 CHMP4B-mNG, mRuby3-NES, CHMP7^NES^*-FLAG cells treated with DOX to induce CHMP7^NES*^-FLAG overexpression. Immunoblot of whole-cell lysates shows CHMP7^NES*^-FLAG levels at indicated time points after DOX wash out. **e**, As in d, quantification of CHMP4B NE foci, CHMP7^NES*^-FLAG mRNA and CHMP7^NES*^-FLAG protein levels before (PRE) DOX induction and following DOX wash out. Bars, SEM. Dots for bar graphs represent the mean of 3 independent experiment; dots for dotplot represent individual measurements, n= 65, 73, 65, 74, 64. ****P<0.0001; ***P<0.001, Dunnett’s multiple comparison test compares time points of each parameter to the corresponding pre-DOX. **f**, As in d, representative confocal images showing CHMP4B-eGFP NE foci. Scale bars, 10 μm. **g**, Overexpression of nuclear CHMP7 drives formation of LEMD2-CHMP4B NE foci. Images of live HeLaK CHMP4B-eGFP, mRuby3-NES and LEMD2-SNAP cells transiently transfected with the indicated CHMP7 alleles and labelled with SiR-SNAP. Nucleus perimeter is outlined. Scale bars, 5 μm.

**Extended Data Figure 2.**
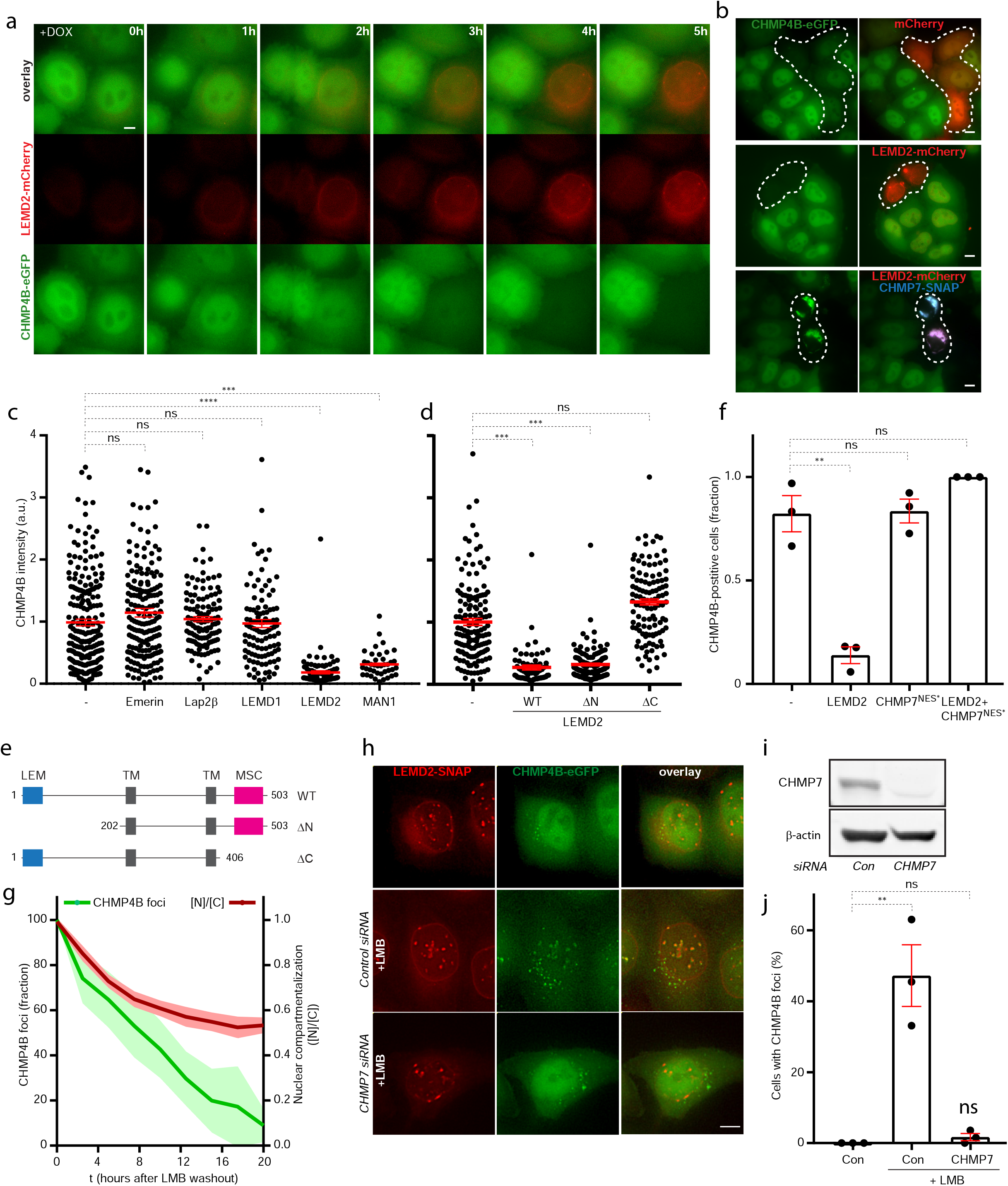
Nuclear localization of CHMP7 stabilizes LEMD2-CHMP7-CHMP4B complexes at the NE. **a**, LEMD2 overexpression selectively depletes CHMP4B levels. HeLaK CHMP4B-eGFP, LEMD2-mCherry cells were treated with DOX to induce overexpression of LEMD2 and were subsequently monitored by live-cell imaging. Scale bar, 5 μm. **b**, LEMD2 overexpression dependent depletion of CHMP4B is countered by co-overexpression of CHMP7. Wide-field images of live HeLaK CHMP4B-eGFP cells transiently transfected with the indicated alleles. Transfected cells are outlined. Scale bar, 5 μm. **c**, LEMD2 overexpression selectively depletes CHMP4B levels. Quantification of CHMP4B intensity in HeLaK CHMP4B-eGFP cells transiently transfected with the indicated mCherry fusion alleles and imaged live. Bars, mean and SEM from 3 experiments. n= 211, 179, 119, 107, 92, 40. ****P<0.0001 and ***P=0.0006 two tailed unpaired *t*-test. df=4. **d**, Quantification of CHMP4B intensity in HeLaK CHMP4B-eGFP cells transfected with mCherry fusions of the indicated LEMD2 alleles and imaged live. Bars, mean and SEM from 3 experiments. n= 145, 67, 166, 111. ***P=0.0001 two-tailed unpaired t-test, df=4. **e**, Schematic representation of LEMD2 functional domains and deletions used in this study. LEM, LAP2, Emerin, MAN1 domain. TM, transmembrane domain; MSC, MAN1-Src1p C-terminal domain. **f**, Interplay between LEMD2, CHMP4B and CHMP7. Quantification of CHMP4B-positive HeLaK CHMP4B-eGFP cells after transfection with LEMD2 and CHMP7 alleles. Bars, mean and SEM from 3 experiments. Dots, mean from each experiment. n= 84, 45, 38, 32. **g**, Interplay between CHMP7 and XPO1 activity. ESCRT-III NE foci induced by LMB treatment were monitored every 30 min by live-cell imaging in HeLaK CHMP4B-eGFP, mRuby3-NES, LEMD2-SNAP cells after LMB wash out. Mean and 95% confidence interval are plotted. n=23 cells from 16 movies. **h**, Formation of CHMP4B NE foci upon XPO1 inactivation was monitored in live HeLaK CHMP4B-eGFP, mRuby3-NES, LEMD2-SNAP cells were treated with indicated siRNAs and LMB where indicated. CHMP4B foci formation depended on CHMP7. Scale bar, 5 μm. **i**, Immunoblot of whole-cell lysate showing depletion of endogenous CHMP7 upon siRNA treatment. **j**, As in h, quantification of cells with CHMP4B NE foci. Bars, mean and SEM from 3 experiments. Dots, mean from each experiment. n= 116, 281, 258. **P=0.0056 two-tailed unpaired *t*-test, df=4.

**Extended Data Figure 3.**
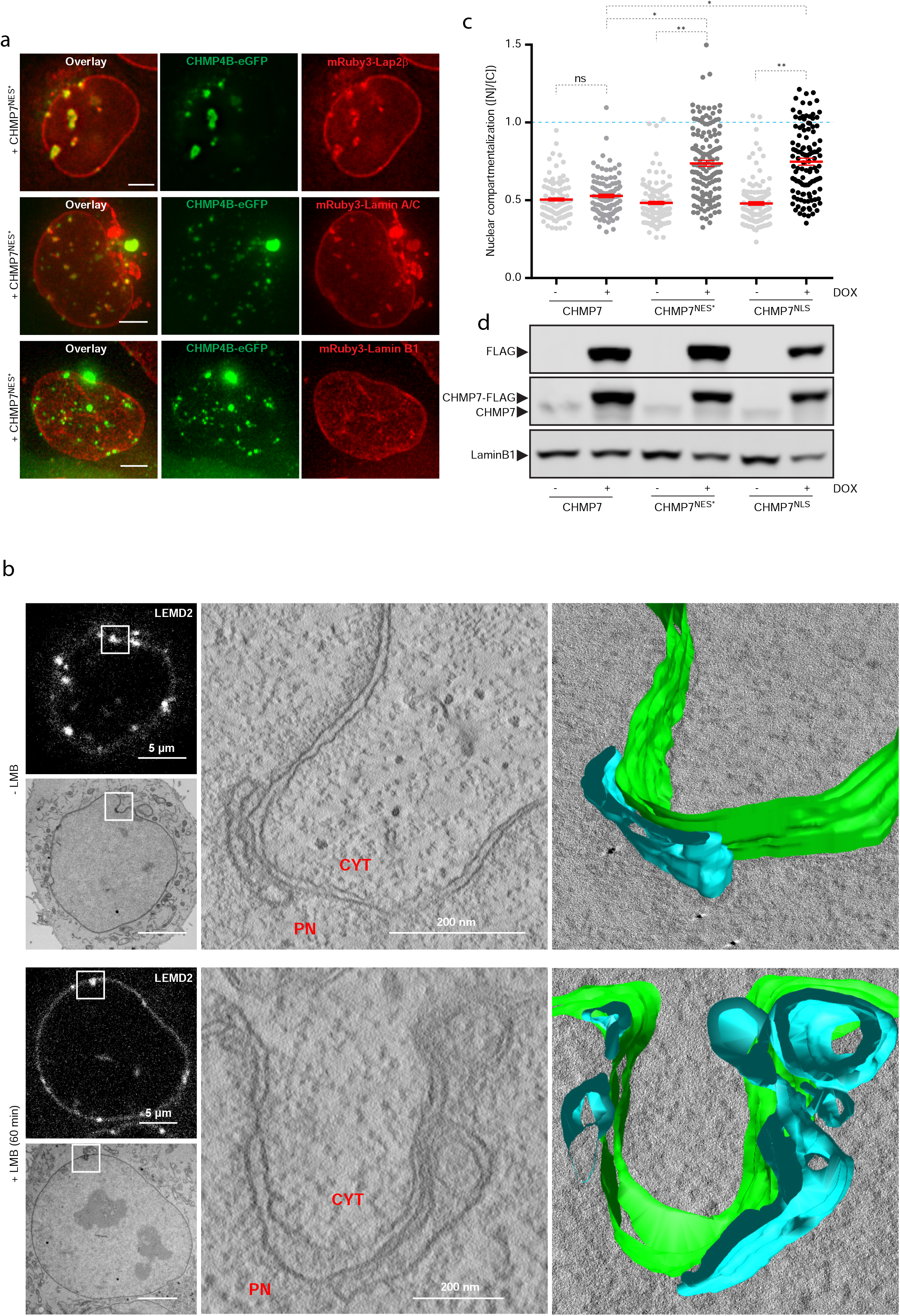
Unrestrained nuclear CHMP7 drives NE deformation and rupture. **a**, Persistent CHMP4B foci colocalize with membrane clusters. HeLaK CHMP4B-eGFP and mCherry-Lap2β, mCherry-Lamin A/C or mCherry-Lamin B1 cells were transiently transfected with CHMP7^NES^*-FLAG and imaged by live-cell imaging. Scale bars, 5 μm. **b**, Inactivation of XPO1 induces 3-dimensional membrane deformations. CLEM from HeLaK CHMP4B-eGFP, mRuby-NES and LEMD2-SNAP cells incubated with LMB where indicated, imaged by live cell imaging, fixed and processed for EM. Slight overexpression of LEMD2-SNAP alone generates LEMD2 foci where the NE displays a parallel sheet of INM. XPO1 inactivation transforms these parallel sheets into a 3D network. Models from tomograms (left panels), Green represents NE, Blue represents INM deformations (top panels). Cyt, cytoplasm; PN, PN. **c**, Overexpression of nuclear CHMP7 results in irreversible membrane rupture. RPE1 CHMP4B-mNG, mRuby3-NES and inducible CHMP7-FLAG, CHMP7^NES^*-FLAG or CHMP7^2NLS^-FLAG cells were treated with DOX to induce the indicated CHMP7 allele and monitored in live-cell imaging. Nuclear compartmentalization was assessed by measuring nucleus/cytoplasm ratio of mRuby3-NES and plotted. Bars, mean and SEM from 3 experiments. n=101, 108, 125, 138, 129, 108. two-tailed *t*-test CHMP7^NES^* DOX-/+ **P= 0.0074, df=4; CHMP7^NLS^ DOX-/+ **P= 0.0054 df=4, CHMP7 DOX+/CHMP7^NES^* DOX+ *P=0.0212 df=4; CHMP7 DOX+/CHMP7^NLS^ DOX+ *P=0.0117. **d**, Immunoblot from whole cell lysates showing expression of endogenous CHMP7 and DOX -induced CHMP7 alleles.

**Extended Data Figure 4.**
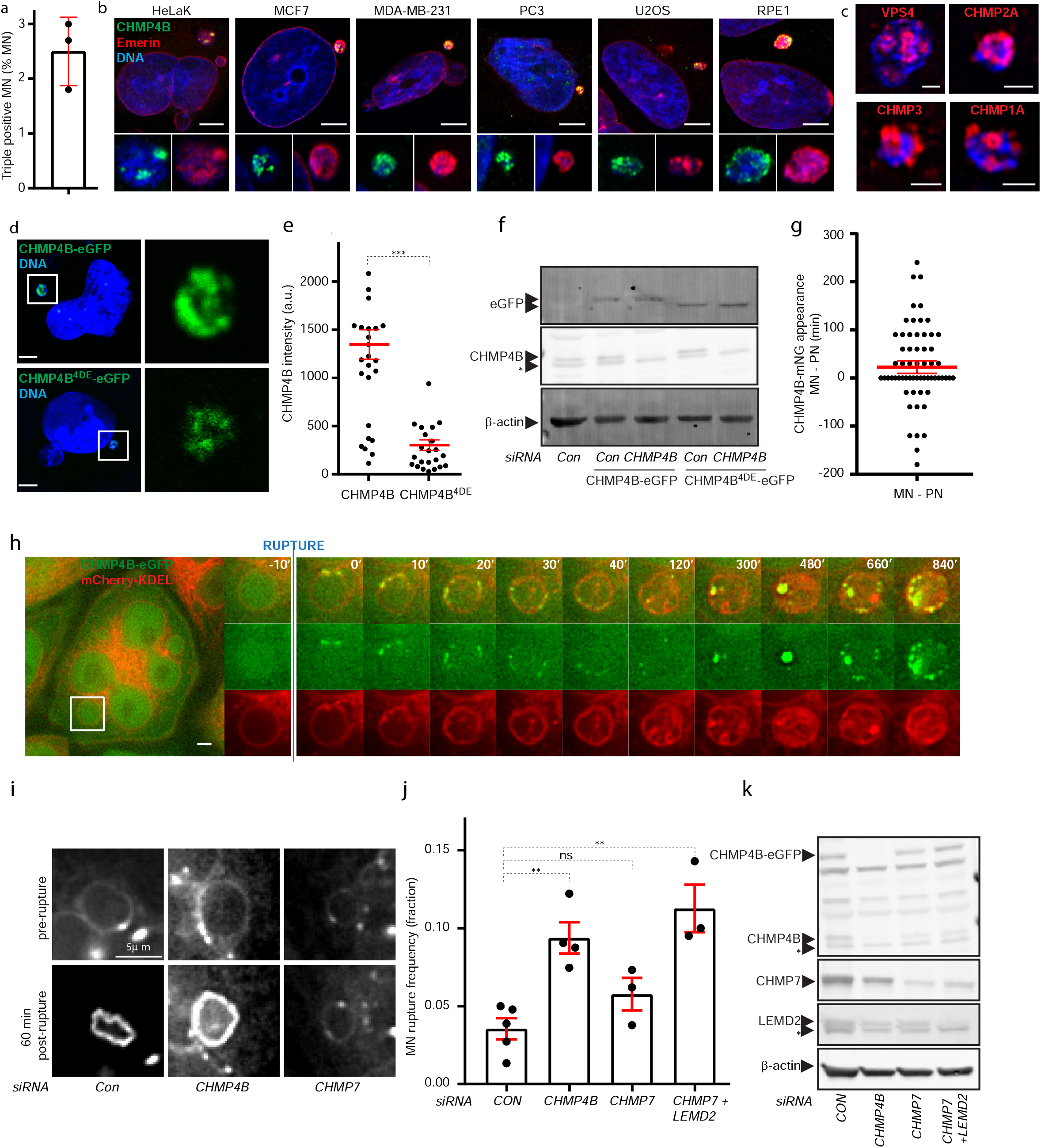
LEMD2, CHMP7 and ESCRT-III are recruited at MN upon rupture. **a**, Quantification of the fraction of MN enriched in CHMP4B-eGFP, endogenous LEMD2 and CHMP7 as in Fig. 3b. Bars, mean and SEM from 3 experiments, n=2666 MN. **b**, CHMP4B localization at ruptured MN in different cell lines. Cells were treated with AZ3146, fixed and stained for endogenous CHMP4B and the NE marker Emerin. Scale bars, 5 μm. **c**, ESCRT-III complex subunits and effectors localize at MN. Confocal images of MN from HeLaK cells fixed and immunolabelled for the indicated endogenous factors. Scale bars, 1 μm. **d**, CHMP4B is recruited to the MNE. Confocal microscopy of HeLaK cells stably expressing siRNA-resistant CHMP4B-eGFP or the membrane-binding defective mutant CHMP4B^4DE^-eGFP, treated with CHMP4B siRNA to deplete endogenous CHMP4B and incubated with AZ3146. Insets show MN. Scale bars, 5 μm. **e**, As in d, quantification of micronuclear GFP fluorescence of the indicated CHMP4B alleles. Bars indicate mean and SEM from 3 experiments. CHMP4B-eGFP n=24; CHMP4B^4DE^-eGFP n=22. ***P=0.0001 two-tailed Mann Whitney Test. **f**, As in d, immunoblot of whole-cell lysate showing depletion of endogenous CHMP4B and expression of the eGFP-fusions. **g**, MN and PN recruit CHMP4B-mNG with similar kinetics upon induction of nuclear CHMP7 (CHMP7^NES^*) expression. Micronucleated RPE1 CHMP4B-mNG, mRuby3-NES and CHMP7^NES^*-FLAG cells were treated with DOX and time of appearance of CHMP4B foci at PN and MN within the same cell was compared by live-cell imaging. n=61. Bars, mean and SEM for 3 experiments. **h**, CHMP4B accumulation and MN collapse. Live-cell imaging of micronucleated HeLaK CHMP4B-eGFP, mCherry-KDEL cells. Gallery shows the evolution of a MN upon rupture. Scale bars, 5 μm. **i**, LEMD2 intensity and MN shape were monitored in live-cell imaging of HeLaK CHMP4B-eGFP, mRuby3-NES and LEMD2-SNAP cells, transfected with indicated siRNA and labelled with SiR-SNAP. Localization of mRuby3-NES was used to determine rupture time. Scale bars, 5 μm. **j**, ESCRT do not induce rupture of MN. Rupture frequency in MN of HeLaK mRuby3-NES cells treated with the indicated siRNAs. Depletion of ESCRT-III or LEMD2 does not rescue rupture of MN. Bars indicate mean and SEM, dots represent the mean of each independent experiment. Con n=821; CHMP4B n=745; CHMP7 n=305; CHMP7+LEMD2 n=256. **k**, Immunoblot from whole-cell lysates showing knockdown efficiency of endogenous proteins as indicated. Asterisks indicate non-specific immunoreactivity.

**Extended Data Figure 5.**
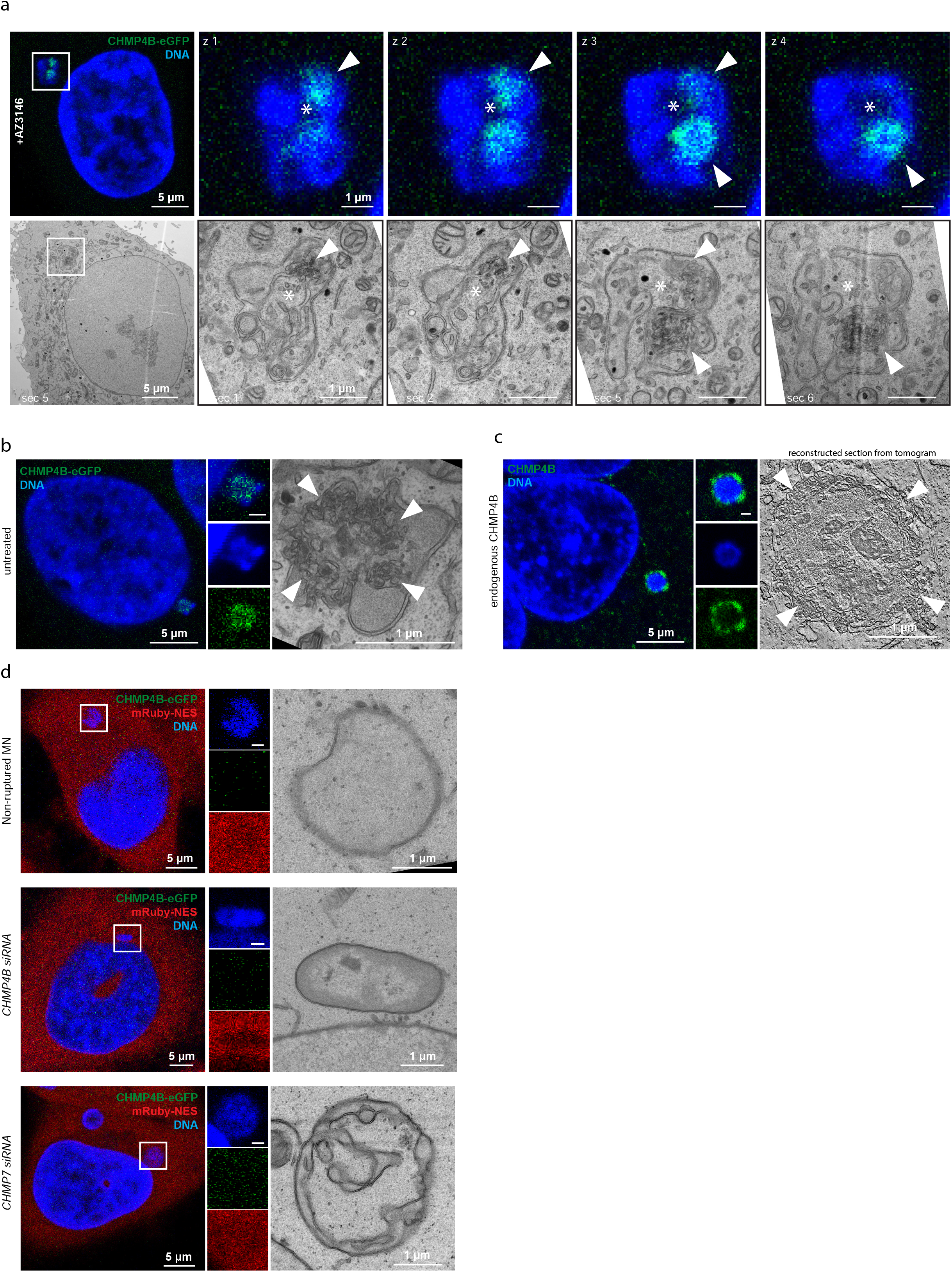
ESCRT-III drives extensive membrane deformations in ruptured MN. **a**, Micronuclear CHMP4B foci correspond to areas of extreme membrane deformations. CLEM of a HeLaK CHMP4B-eGFP cell, treated with AZ3146 to induce MN, fixed, imaged with confocal microscopy and then processed for EM. Position of CHMP4B foci (detected by light microscopy) relative to areas of the MN where the NE is dramatically deformed (observed on consecutive EM sections). Subsequent confocal planes with the respective electron micrographs are shown. Arrowheads indicate confocal planes of CHMP4B-eGFP foci and the corresponding areas in the electron micrographs. Asterisks mark are included as reference for MN position. **b**, CHMP4B foci (detected by light microscopy) correlate with MNE deformations (arrowheads) in naturally occurring MN (observed on EM sections, arrowheads) of HeLaK CHMP4B-eGFP cells. **c**, Endogenous CHMP4B foci (detected by light microscopy) correlate with MNE deformations (arrowheads) in MN (observed on tomograms from consecutive EM sections) of HeLaK cells. **d**, ESCRT-III depletion rescues MNE trabecular distortions. CLEM of ruptured MN in HeLaK CHMP4B-eGFP, mRuby-NES cells transfected with indicated siRNAs, incubated with AZ3146, fixed, imaged with light microscopy and processed for EM.

**Extended Data Figure 6.**
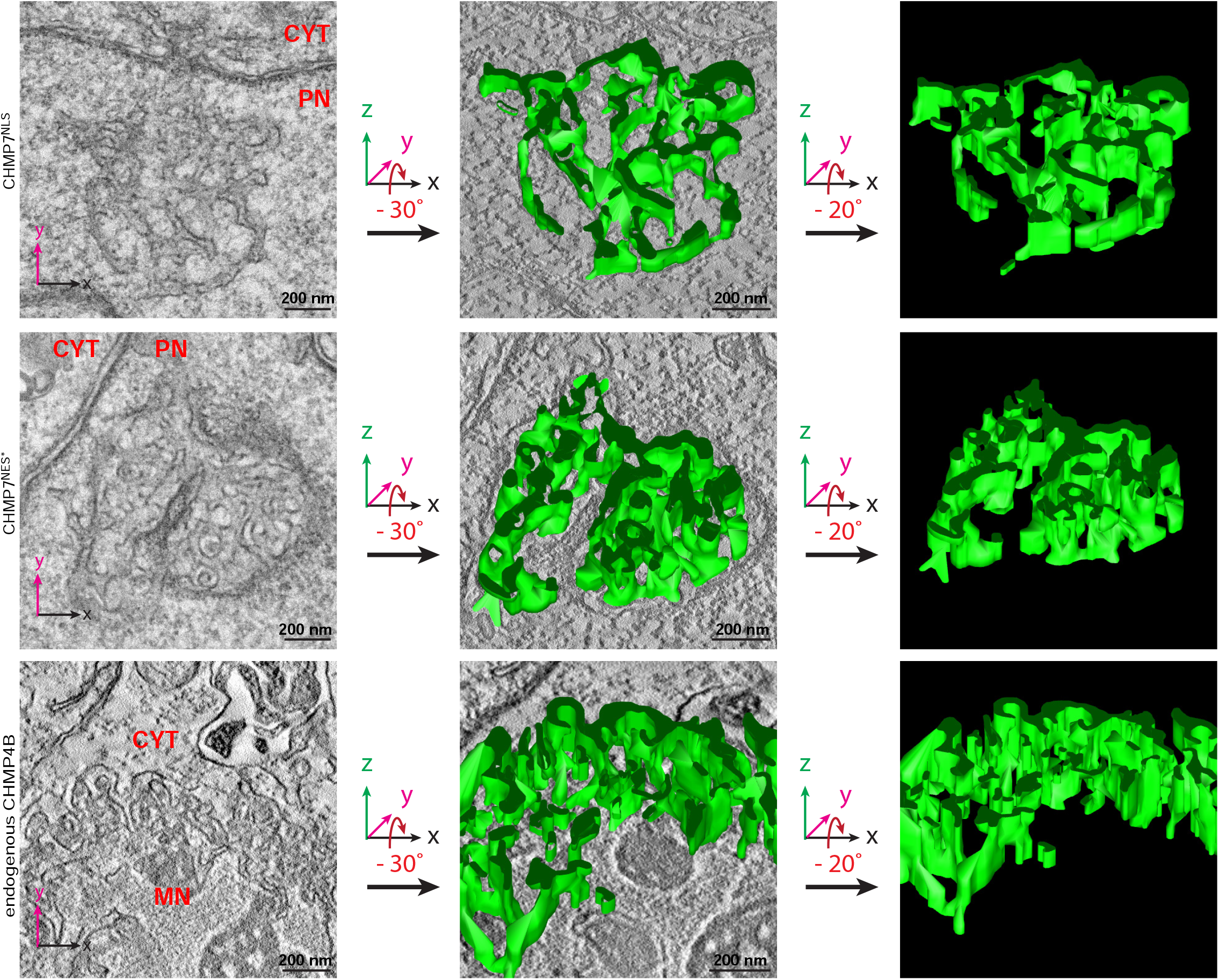
Ruptured MN ultrastructure resemble trabecular membrane network driven by CHMP7 overexpression. Electron tomograms from CLEM experiments and 3D-models of membrane ultrastructure induced by overexpression of CHMP7^NLS^-FLAG (upper panels) or CHMP7^NES^*-FLAG (middle panels). Note the similarity to a “spontaneously” ruptured MN of HeLaK cells with distinct endogenous CHMP4B foci (from Extended Data Fig. 5c) (lower panels).Cyt, cytoplasm; PN, primary nucleus; MN, micronucleus.

**Extended Data Figure 7.**
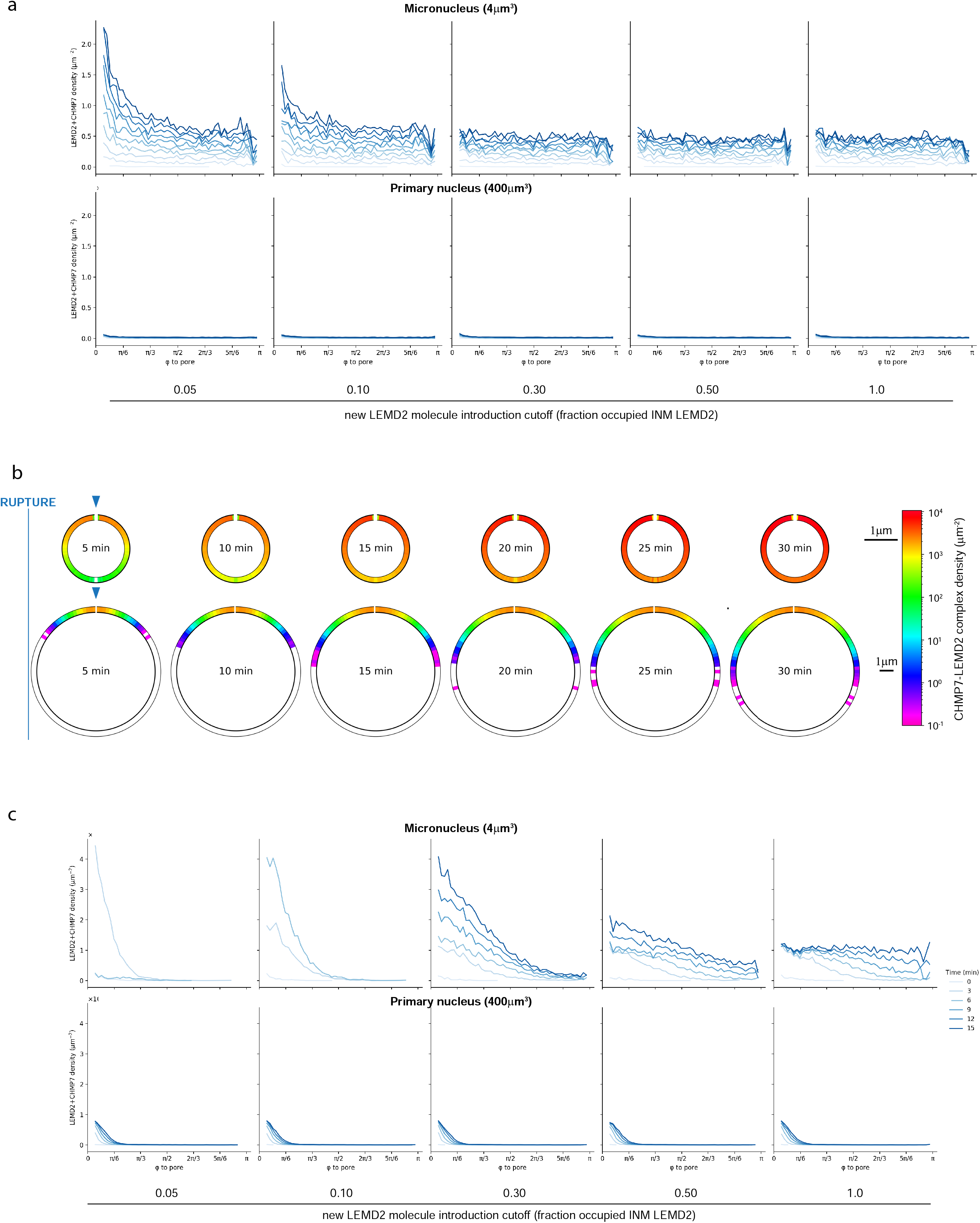
MN are unable to restrict CHMP7-LEMD2 complexes. **a**, Phi line plots from *in silico* experiments showing evolution of CHMP7-LEMD2 complex angular density along the INM from the rupture site (φ to pore = 0) to the opposite pole (φ to pore =π) for cytoplasmic CHMP7. Results are shown for micronuclei (top panel) and primary nuclei (bottom panel) at different time points after rupture (as indicated), and using different LEMD2 occupancy cut-offs for generation of new INM LEMD2 molecules. **b**, as Fig. 3H, but with CHMP7 as an ER membrane-bound protein. **c**, as a, but with CHMP7 as an ER membrane-bound protein.

**Extended Data Figure 8.**
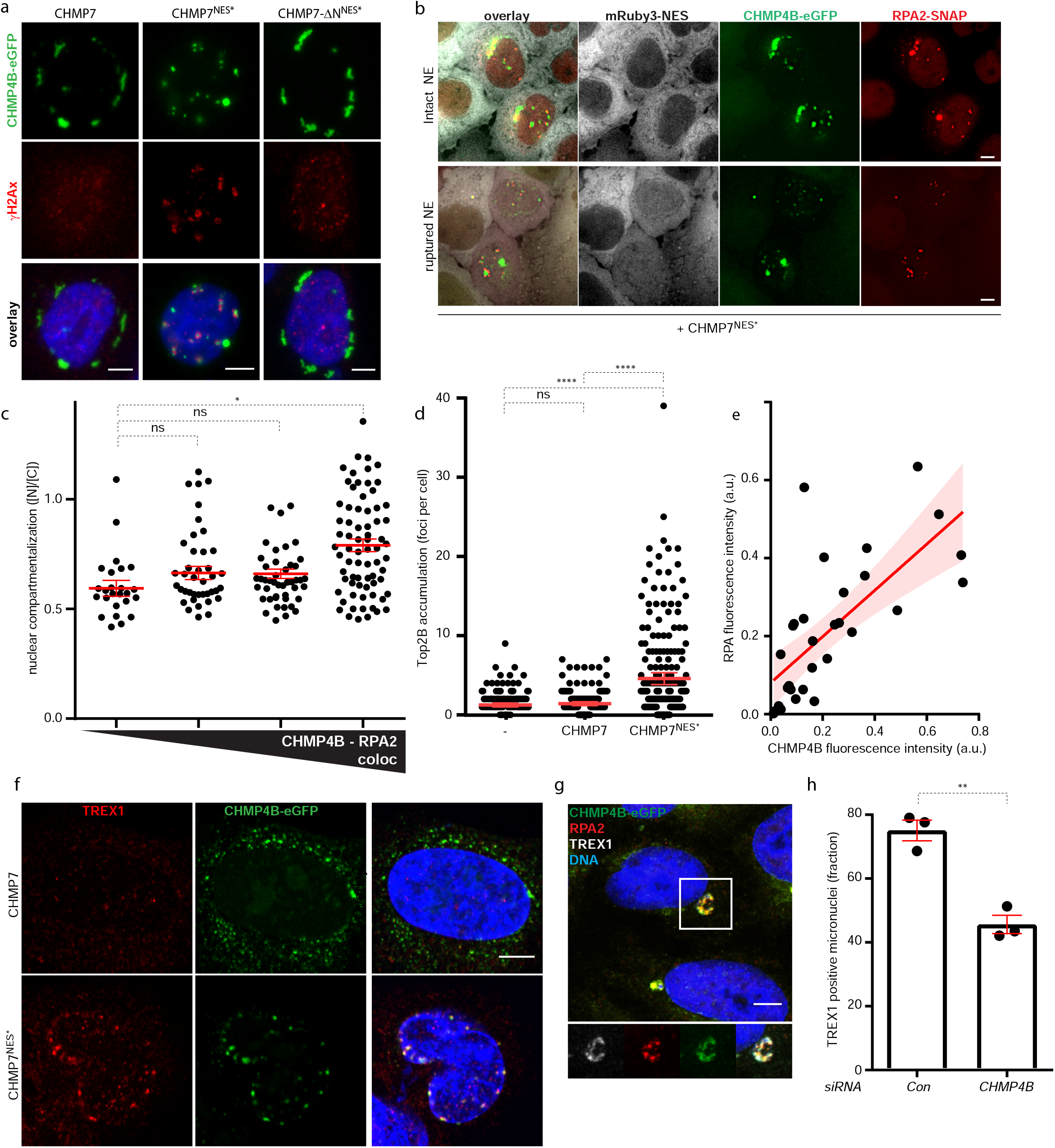
Unrestrained nuclear ESCRT-III and its consequences for the genome. **a**, Nuclear CHMP4B foci associate with local DNA damage. RPE1 CHMP4B-mNG, mRuby3-NES and inducible CHMP7-FLAG, CHMP7^NES^*-FLAG or CHMP7-ΔN^NES^*-FLAG (unable to bind to membranes) were treated with DOX, fixed and stained for the DNA damage marker γH2Ax. Scale bars, 5 μm. **b**, RPA2 accumulates at CHMP4B foci independently of NE ruptures. Wide-field images of live HeLaK CHMP4B-eGFP, RPA-SNAP, mRuby-NES cells transiently overexpressing CHMP7^NES^*-FLAG and stained with SiR-SNAP. Scale bars, 5 μm. **c**, Quantification from experiment described in b. Nuclear leakiness was measured by nucleus/cytoplasm ratio of mRuby-NES intensity and increasing degrees of CHMP4B-RPA2 colocalization were plotted. n= 25, 40, 45, 72. *P=0.014 two-tailed unpaired *t*-test, df=4. **d**, Nuclear CHMP7 induces DNA torsional stress. RPE1 CHMP4B-mNG, mRuby3-NES and inducible CHMP7-FLAG or CHMP7^NES^*-FLAG cells were treated with DOX to induce the indicated CHMP7 allele, fixed, stained for Top2B and imaged by confocal microscopy. The number of nuclear foci of TOP2B was quantified. Representative images are shown in Fig.4b. Bars, mean and SEM. Control (-) n=255; CHMP7 n= 136; CHMP7^NES^*= 205. ****P < 0.0001 two-tailed Mann Whitney Test. **e**, CHMP4B intensity at MN correlates with RPA accumulation. Fixed HeLaK cells were immunolabelled for CHMP4B and RPA2. RPA2 and CHMP4B fluorescence intensities were measured within the same MN and plotted on the x- and y-axes, respectively. Regression line (line) and 95% confidence interval (bands) are indicated. R^2^= 0.4978, n= 31. **f**, The ER-associated TREX1 exonuclease enriches at nuclear CHMP4B foci. RPE1 CHMP4B-mNG, mRuby3-NES and inducible CHMP7-FLAG or CHMP7^NES^*-FLAG cells were treated with DOX to induce the indicated CHMP7 allele, fixed, stained for TREX1 and imaged by confocal microscopy. Scale bars, 5 μm. **g**, TREX1 is enriched at CHMP4B foci in collapsed MN. CHMP4B-eGFP, mRuby3-NES, RPA2-SNAP HeLaK cells were treated with AZ3146 to induce MN, fixed and stained for RPA2 and TREX1. Scale bar, 5 μm. **h**, Quantification of ruptured MN showing TREX1 enrichment. mRuby3-NES, CHMP4B-eGFP HeLaK cells were treated with Control siRNA or a CHMP4B siRNA targeting both endogenous and GFP-tagged allele. Cells were then treated with AZ3146 to induce formation of MN, fixed and stained for TREX1. Bars indicate mean and SEM, dots represent the mean of each independent experiment. Con, n=171; CHMP4B, n=238. **P=0.0025 two-tailed unpaired *t*-test, df=4.

**Extended Data Figure 9.**
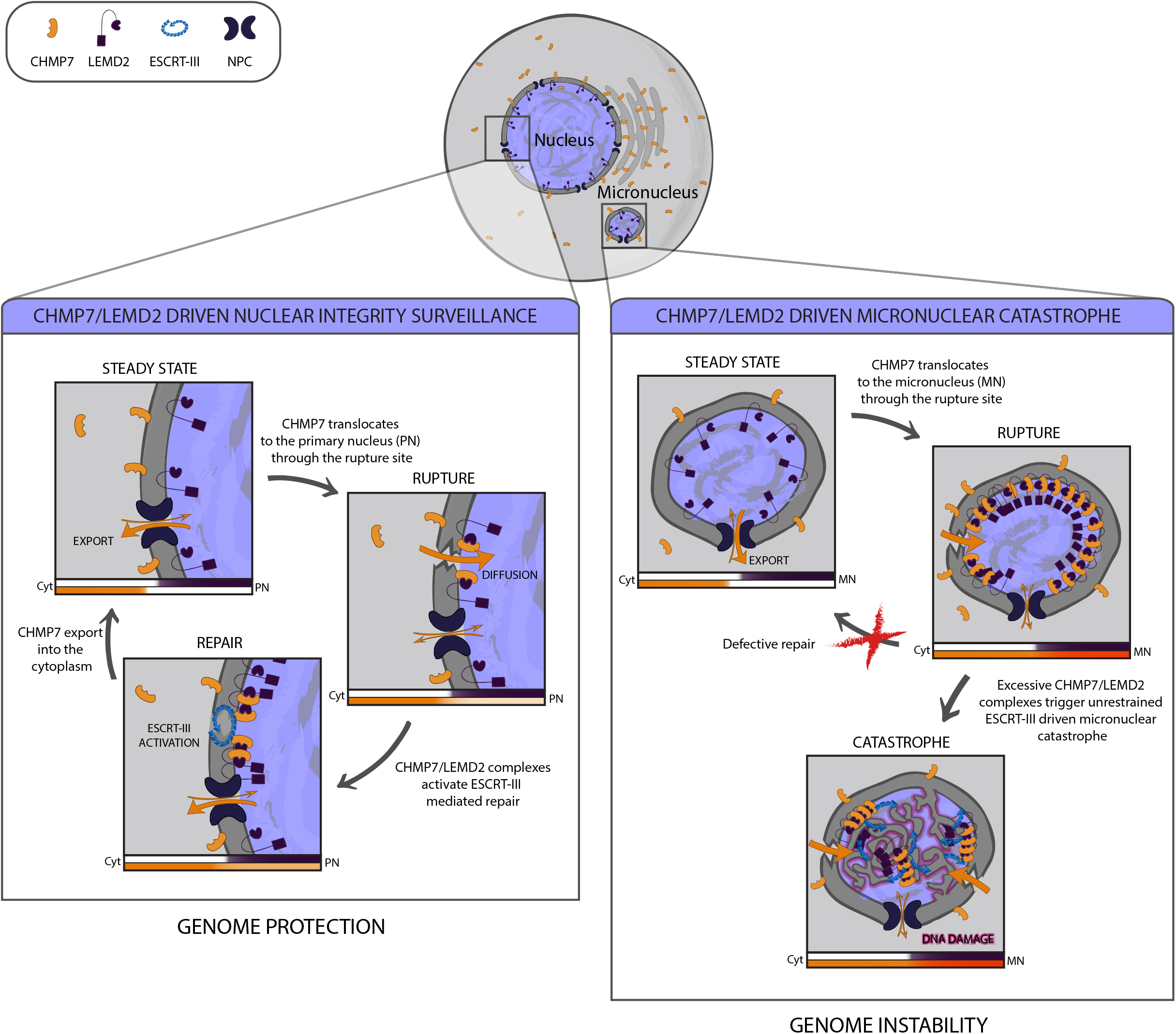
Model for ESCRT-III function at NE of primary nuclei and micronuclei. When the NE is intact (steady state) CHMP7 is continuously exported from the nucleus in an XPO1-dependent fashion, precluding its interaction with LEMD2 at the INM. Upon rupture, CHMP7 freely diffuses into the nucleus through the site of rupture. LEMD2-CHMP7 complexes form in proximity of the rupture site to activate ESCRT-III polymerization and sealing activity (repair). MN are intrinsically unable to restrain ESCRT-III activity. Upon rupture, CHMP7 diffuses into the MN, quickly spreads along the whole INM surface and rapidly saturates all available LEMD2. This lack of CHMP7-LEMD2 restriction causes unrestrained ESCRT-III activity that is unable to repair the membrane and instead drives extreme membrane deformation, MN catastrophe and DNA damage. The bar at the bottom of each illustration represents the cytosolic vs (micro)nuclear distribution of CHMP7.

## Methods

### Cell cultures

Cell lines were cultured as recommended by ATCC guidelines. For live-cell imaging, cells were seeded into glass bottom imaging dishes (Lab-Tek, Nunc; MatTek, MatTek Corporation). For immunofluorescence studies, cells were grown on high Precision cover glass (Marienfeld). For confocal microscopy, cells were fixed with 4% EM grade Formaldehyde (Polysciences) for 5 minutes at 37 °C. Cells were then permeabilized by treatment with 0.16% Triton X-100 for 2 minutes. Primary and secondary antibodies were diluted in PBS/0.05% Saponin and incubated for 1 to 2 hours. After antibody staining, samples were mounted on microscope slides (Menzel-Glaser) with Mowiol for standard confocal imaging or SlowFade Gold Antifade Reagent (Life Technologies) for Airyscan microscopy. RPE1 and HeLaK cell lines stably expressing inducible CHMP7 or LEMD2 alleles were treated with 500 ng/ml doxycycline (DOX, Sigma-Aldrich). XPO1 inhibition was carried on by incubating cells with 2.5 ng/ml Leptomycin B (LMB, InvivoGen) for up to 3 hours. Loss of XPO1 activity was monitored my nuclear localization of 2xmRuby3-NES. Micronucleation was induced by MPS1 inhibitor AZ3146 (Selleckcem) used at a concentration of 2 μM. RPE1 cell lines were depleted for p53 before AZ3146 treatment.

### Stable cell lines

A stable HeLa “Kyoto” cell line expressing CHMP4B-eGFP was obtained from Anthony Hyman^1^. All other stable cell lines were lentivirus-generated pools. To achieve low expression levels, the weak PGK promoter was used for transgene expression. For higher expression levels CMV or EF1α promoters were used. Third generation Lentivirus was generated using procedures and plasmids as previously described^2^. Briefly, (eGFP/mNG/mCherry/SNAP/FLAG fusions of) transgenes were generated as Gateway ENTRY plasmids using standard molecular biology techniques, with all CHMP4B-fusions containing the LAPtag module^1^. From these vectors, Lentiviral transfer vectors were generated by recombination into lentiviral destination vectors (Addgene plasmids # 19067, 19068, 41393; and vectors derived from pCDH-EF1α-MCS-IRES-PURO (SystemBiosciences, inc.)) using Gateway LR reactions. VSV-G pseudotyped lentiviral particles were packaged using a third generation packaging system^3^ (Addgene plasmids # 12251, 12253, 12259). HeLaK or RPE1 cells were then transduced with low virus titers (MOI ≤1) and stable expressing populations were generated by antibiotic selection. Detailed cloning procedures are available on request. The stable cell lines used in this study are listed in Supplementary Data 1.

### Transient plasmid transfections

For transient expression of CHMP7 and LEMD2 alleles, cells were grown on Precision cover glass in 24-well plates or into glass bottom imaging dishes and transfected with 0.5 μg DNA construct complexed with Fugene6 (Promega) following the manufacturer’s instruction. Cell were then fixed or imaged by live-cell imaging 12 to 24 hours after transfection. Non-targeting control Silencer Select siRNA (pre-designed, Cat. No. 4390844) was used as control. The plasmids used in this study are listed in Supplementary Data 1, and detailed cloning procedures are available on request.

### siRNA transfections

All siRNAs were purchased from Ambion and contained the Silencer Select modification. Cells at 50% confluency were transfected with Lipofectamine RNAiMAX transfection reagent (Life Technologies) following the manufacturer’s instruction. Cells were transfected with 50 nM CHMP2A siRNA (AAGAUGAAGAGGAGAGUGAtt) or p53 (GUAAUCUACUGGGACGGAAtt) for 48 hours. CHMP4B (CAUCGAGUUCCAGCGGGAGtt), CHMP7 (AGGUCUCUCCAGUCAAUGAtt) and LEMD2 (AGCUGGUAAUUUUGAGUGUtt) were silenced by 2x serial transfections with 50 nM siRNA concentration for a total of 72 hours.

### Antibodies

Rabbit anti-human CHMP4B antibody (WB 1:2000, IF 1:1000) and rabbit anti-CHMP3 (IF 1:1000) were produced as previously described^4,5^. Rabbit anti-CHMP2A (Proteintech #10477-1-AP; WB 1:500, IF 1:100), rabbit anti-CHMP7 (Proteintech #16424-1-AP; Sigma-Aldrich; WB 1:250), mouse anti-CHMP7 (Abnova #H00091782-B01; IF 1:100), rabbit anti-LEMD2 (Abnova #PAB20940; WB 1:500; IF 1:100), mouse anti-CHMP1A (Abcam, #ab104100; IF 1:150), rabbit anti-VPS4 (Sigma-Aldrich #SAB4200025; WB 1:500, IF 1:200), mouse anti-γH2Ax (Millipore 05-636; IF 1:100), mouse anti-Emerin (NeoMarkers MS-1751; IF 1:100), mouse anti-RPA2 (Abcam #ab2175; IF 1:100), rabbit anti-TREX1 (Abcam #ab185228; IF 1:200), mouse anti-TOP2B (scbt #sc-25330; IF 1:200), mouse anti-β-Actin (Sigma-Aldrich; WB 1:30000), mouse anti-GFP (Roche #1814460001; WB 1:1000; IF 1:200), rabbit anti-FLAG (Cell Signaling Technology 2368; WB 1:1000; IF 1:400), rabbit anti-Histone H3 (Abcam #ab1791; WB) and mouse anti-Lamin B1 (scbt #sc365962; WB 1:1000) were used as primary antibodies. DNA was labelled with Hoechst 33342 (2 μg/ml, Life Technologies. Secondary antibodies included anti-mouse-, anti-rabbit- and anti-goat-Alexa488 (Jackson), Alexa568 (molecular Probes) and Alexa647 (Jackson).

### Confocal microscopy

Fixed samples were imaged with a Zeiss LSM 710 or 780 confocal microscope (Carl Zeiss MicroImaging GmbH, Jena, Germany) equipped with an Ar-Laser Multiline (458/488/514 nm), a DPSS-561 10 (561 nm), a Laser diode 405-30 CW (405 nm), and a HeNe-laser (633 nm). The objective used was a Zeiss Plan-Apochromat 63x/1.40 Oil DIC M27. Image processing was performed with basic software ZEN 2009 (Carl Zeiss MicroImaging GmbH, Jena, Germany) and ImageJ software^6^ (National Institutes of Health, Bethesda, MD, USA).

### Confocal Airyscan microscopy

Fixed cells were imaged with a Zeiss LSM 880 Airyscan microscope (Carl Zeiss MicroImaging GmbH, Jena, Germany), equipped with an Ar-Laser Multiline (458/488/514nm), a DPSS-561 10 (561nm), a Laser diode 405-30 CW (405 nm), and a HeNe-laser (633nm). The objective used was a Zeiss plan-Apochromat 63xNA/1.4 oil DICII. The images were acquired with the Airyscan detector with a voxel size of 0.035 × 0.035 × 0.144μm for high resolution imaging and super-resolution processing resulting in a final resolution of 120 × 120 × 350 nm. Image acquisition, processing and analysis were performed with the ZEN 2.3 SP1 basic software (Carl Zeiss), or with ZEN 2.3 Blue (Carl Zeiss). 3D visualization of z-stacks was done using Imaris 7.7.2 (Bitplane AG, Zürich, Switzerland). Manders’ and Pearson’s colocalization measurements were done in ZEN 2.3 Blue with a Costes automatic thresholding. Bitplane Imaris software was used to generate surface renderings of 3D-SIM images.

### Live microscopy

Cells seeded into glass bottom imaging dishes were imaged on a DeltaVision microscope (Applied Precision) equipped with Elite TruLight Illumination System, a CoolSNAP HQ2 camera and a 40× or 60× Plan Apochromat (1.42 NA) lenses. For temperature control during live observation, the microscope stage was kept at 37 °C by a temperature-controlled incubation chamber. Time-lapse images deconvolved using the softWoRx software (Applied Precision, GE Healthcare) and processed with ImageJ for presentation and quantifications. Cells were imaged in DMEM gfp-2 anti-bleaching live-cell imaging medium (Evrogen) supplemented with 10% fetal bovine serum (FBS). Environmental control was provided by a heated stage and an objective heater (20-20 Technologies). SNAP-Cell^®^ 647-SiR (SiR-SNAP 1:2000) (New England Biolabs) was used to detect SNAP-tagged alleles. SiR-DNA (1:2000) (SpiroChrome)^7^ was used to monitor nuclei and MN. Images were deconvolved using softWoRx software and processed in ImageJ/FIJI^6^ or Imaris. Nuclear ruptures are identified by compromised nucleocytoplasmic compartmentalization as visualized by the equalization of 2xmRuby3-NES intensity between the (micro-)nucleus and cytoplasm. For quantification of nucleo-cytoplasmic compartmentalization in living cells (Fig.2a, Extended Data Fig.3c, 8c), regions of interest (ROIs) were drawn in the nucleus and cytoplasm, followed by extraction of the mean fluorescence intensity, background fluorescence subtraction, and calculation of [N]/[C] ratios. For cells with intact nuclei [N]/[C] ratio approximates 0.5, while for cells with ruptured nuclei this ratio approximates 1.

### High-content microscopy

For quantification of LEMD2-CHMP7-CHMP4B positive MN and TREX1 enrichment experiments Olympus ScanR system (illumination system with an UPLSAPO 40X objective) was used for imaging of a large number of cells in fixed samples and stained for the proteins of interest.

### Correlative light and electron microscopy

Cells for CLEM experiments were grown on photo-etched coverslips (Electron Microscopy Sciences, Hatfield, USA). Fixation was done with 4% formaldehyde, 0.1% glutaraldehyde in 0.1 M PHEM (240 mM PIPES, 100 mM HEPES, 8 mM MgCl2, 40 mM EGTA), pH 6.9, for 1h. The coverslips were washed with 0.1 M PHEM buffer and mounted with Mowiol containing 1 μg/ml Hoechst 33342. The cells were examined with a Zeiss LSM710 confocal microscope (Carl Zeiss MicroImaging GmbH, Jena, Germany) equipped with a Laser diode 405-30 CW (405nm), an Ar-Laser Multiline (458, 488, 514 nm), a DPSS-561 10 (561 nm) and a HeNe-laser (633 nm). Cells of interest were identified by fluorescence microscopy and a Z-stack covering the whole cell volume was acquired. The relative positioning of the cells on the photo-etched coverslips was determined by taking a low magnification DIC image. The coverslips were removed from the object glass, washed with 0.1 M PHEM buffer and fixed in 2% glutaraldehyde/0.1 M PHEM overnight. Cells were postfixed in osmium tetroxide and potassium ferry cyanide, stained with tannic acid, and uranyl acetate and thereafter dehydrated stepwise to 100% ethanol followed by flat-embedding in Epon. Serial sections (80 nm – 200 nm) were cut on a Ultracut UCT ultramicrotome (Leica, Germany) and collected on formvar coated slot-grids. Thin sections were observed at 80 kV in a JEOL-JEM 1230 electron microscope and images were recorded using iTEM software with a Morada camera (Olympus, Germany). Sections of 200 nm thickness were observed at 200 kV in a Thermo ScientificTM TalosTM F200C microscope and recorded with a Ceta 16M camera. Consecutive sections were used to align electron micrographs with fluorescent images in X, Y and Z. For tomograms, image series of 200 nm thick section were taken between −60° and 60° tilt angles with 2° increment. Single-tilt axes series were recorded with a Ceta 16M camera. Tomograms were computed using weighted back projection using the IMOD package. Display, segmentation and animation of tomograms were also performed using IMOD software version 4.9^8^.

### Immunoblotting

Cells were lysed in 2x sample buffer (100 mM Tris-HCl pH 6.8, 4% SDS, 20% glycerol, 200 mM DTT, bromophenol blue). The whole-cell lysate was subjected to SDS-PAGE on a 4-20% gradient gel. Proteins were transferred to Immobilon-P membrane (Millipore). Immunodetection was performed using fluorescently-labeled secondary antibodies and Odyssey^®^ or ChemiDoc^™^ developer. Western blot images are representatives for at least three independent experiments.

### Real-Time quantitative PCR (RT-qPCR) analysis

Total RNAs were extracted using RNeasy^®^ Mini Kit, ref. 74106, Qiagen, Germany. Reverse transcription was performed on 500 ng of DNAseI treated total RNA, with high-capacity cDNA reverse transcription kit, following manufacturer’s recommendations (Applied Biosystems, ref. 4368814, USA). RT-qPCR was performed on 2 μl of 1/25 diluted cDNA with a CFX-96 (Biorad) thermocycler, using SsoAdvanced™ Universal SYBR Green Supermix 2X, Biorad. The following primers were used (Invitrogen): *CHMP7* (Forward GTCACAGTCCTCGAGCAGAA / Reverse CGTCATTGACTGGAGAGACCT), *CHMP7-FLAG* (Forward GTCACAGTCCTCGAGCAGAA / Reverse TACGTCGTTAACAGGGCTCA), *GAPDH* (Forward TCGGAGTCAACGGATTTGGT / Reverse TATGATTCCACCCATGGCAA) at a final concentration of 250 nM. All measurements were normalized to the expression of *GAPDH*. Results were analysed using the 2^−ΔΔCT^ method and are presented as the mean of three independent experiments.

### Metaphase spreads

RPE1 cells stably expressing 2xmRuby-NES and DOX-inducible CHMP7-FLAG or CHMP7NES*-FLAG were treated with 500 ng/ml DOX. After 4 hours, colchicine was added to the medium, the cells were incubated for another 20 hours, and harvested according to Mandahl (1992). Chromosomes of the dividing cells were then G-banded with Wright stain (Merck, Darmstad, Germany). The subsequent cytogenetic analysis and karyotypic description followed the recommendations of the International System for Human Cytogenetic Nomenclature (ISCN 2106).

### *In silico* experiments

The geometry of the *in silico* experiments consists of a spherical cell and a smaller, concentric nucleus, populated by two types of particle, CHMP7 and LEMD2. LEMD2 is located on the INM and CHMP7 is either located in the cytoplasm or on the outer nuclear membrane (ONM)/ER. During the *in silico* experiments, the number of the LEMD2 sites bound with CHMP7 is monitored following simulated rupture of the nuclear envelope. The interaction is a reversible reaction with a forward rate of *F_L_* and a backwards rate *D_L_*. The values for all relevant variables are listed in Supplementary Table 1.

**Supplementary Table 1.** Variables used for *in silico* experiments.

| Supplementary Table 1. Variables used for *in silico* experiments. |  |
| --- | --- |
| Parameter | Value |
| CHMP7+LEMD2 binding affinity | 40 nM |
| Cell volume | 2000 $\mu\text{m}^3$ |
| Primary nucleus volume | 400 $\mu\text{m}^3$ |
| Micronucleus volume | 4 $\mu\text{m}^3$ |
| Rupture diameter | 100 nm |
| CHMP7 density (Cytoplasmic)<sup>9-13</sup> | 12 /$\mu\text{m}^3$ |
| CHMP7 density (ER-associated)<sup>9-13</sup> | 2 /$\mu\text{m}^2$ |
| LEMD2 density<sup>10-13</sup> | 100 /$\mu\text{m}^2$ |
| CHMP7 diffusion rate (Cytoplasmic)<sup>14,15</sup> | 40 $\mu\text{m}^2/\text{s}$ |
| CHMP7 diffusion rate (ER-associated)<sup>16,17</sup> | 0.5 $\mu\text{m}^2/\text{s}$ |
| LEMD2 diffusion rate<sup>18,19</sup> | 0 $\mu\text{m}^2/\text{s}$ |
| Bound LEMD2 limit | 0.5 |

For localization, two settings for CHMP7 are considered. In the first case, CHMP7 is treated as soluble and is free to diffuse in the cytoplasm in 3D. In the second case, CHMP7 is tethered to the ONM/ER and diffusion is restricted to 2D on the surface. Since the surface area of the ER is orders of magnitude larger than the nuclear membrane, we assume the CHMP7 particles on the ONM are in contact with a reservoir: when a particle moves from the outer to inner membrane, a new particle is placed randomly on the ONM in order to keep the concentration constant.

Three phenomena were modeled to capture the full behaviour of the system: diffusion, reactions and decay. In free (3D) diffusion, a particle is expected to move a distance 〈*δx*〉 after a time of *δt* as given by 〈*δx*^2^〉 = 6*Dδt*, where *D* is the diffusion constant of the particle. In our model we implement this by allowing each particle to jump a distance, *j*, in 3D space at each time-step with a probability, *P_walk_*, given by

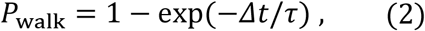

where *τ* is the waiting time of the particle and *Δt* is the time-step of the simulation. The value of *τ* is chosen to ensure consistency between our simulation and the expected physical results:

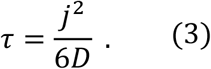

This particular implementation was chosen to simplify the calculation of reaction probabilities.

In the soluble (3D) CHMP7 scenario, if the jump would take the particle through a membrane barrier, the particle is bounced off the membrane instead, unless it would pass through the nuclear envelope rupture. Once a CHMP7 particle is bound to a LEMD2 particle it is immobilized until the complex decays.

In the case of membrane-bound CHMP7 (2D), diffusion is modeled on a spherical surface. The particles were created on the sphere by picking their location uniformly for *θ* ∈ [0,2*π*) and cos*φ* ∈[−1,1), since picking the positions uniformly in *φ* would result in an overrepresentation of particles at the poles^20^. To ensure that a particle moves in a uniform manner, a temporary particle *P*_0_ was generated on the sphere a distance *j* away from the north pole at an angle *ψ* ∈ [0,2*π*). The particle is then rotated twice about the origin of the sphere, using the same rotations that would carry the pole of the sphere on to the old position of the particle at (*θ*, *φ*). As distances are conserved during the rotation, the new particle is now a distance j away from its old position at a uniformly distributed angle, as we required. This means the resultant position *P* is given by;

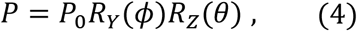

where *R_Y_* and *R_Z_* are the standard 3D rotation matrices.

Following a CHMP7 particle jump, the distance to the closest LEMD2 particle is measured. If this distance is within a collision distance, *r*_c_, the probability for a CHMP7 particle to combine with a LEMD2 particle and act as a single complex until decay is given by

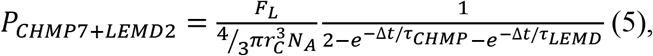

where *N_A_* is the Avogadro’s constant.

The full derivation of this is given in ^21^. From eq. 5, we are free to choose the value of *r_c_* to fix *P_CHMP7+LEMD2_* (or the probity to fix the radius) Decreasing the collision radius increases the accuracy of the simulation, provided we do not allow *P* > 1. To prevent exceeding this limit, a constant value of *r_c_* = 50 nm for both reactions is chosen.

As the reaction are bi-directional, separation of the complexes was also considered. Provided that the decay rate is much lower than the frequency of the simulation time steps, the probability of a CHMP7·LEMD2 complex dissociating into its separate component particles per time-step is defined as,

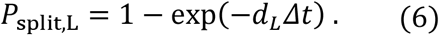

Data visualization focused on the density of the LEMD2-bound CHMP7 particles and the location of these complexes.

### Image processing

For measurement of LEMD2 intensity increase and changes in MN circularity upon rupture (Fig. 3e,f), the perimeter of the MN was manually drawn using LEMD2 signal as membrane proxy. The z-section corresponding to the middle plane of each MN was chosen. For quantification of CHMP4B-eGFP and CHMP4B^4DE^-eGFP intensity at MN (Extended Data Fig. 4d), MN area was segmented by Otzu thresholding based on the DNA channel. The ImageJ function “Analyze Particles” was then used to define ROIs and the mean intensity for the GFP channel of each ROI was measured. A band-shaped ROI around MN was used to measure mean intensity of the cytoplasm as regularization factor. Image processing was performed using ImageJ/FIJI software^6^. For quantification of TOP2B accumulation (Extended Data Fig. 8d), the area around nuclei was selected according to DNA staining and number of TOP2B foci was measured by using the “Count Maxima” function with tolerance values determined for each experiment depending on the imaging parameters. For measurement of CHMP4B and RPA2 fluorescence intensity at MN (Extended Data Fig. 8e), MN area was segmented by Otzu thresholding based on the DNA channel. The ImageJ function “Analyze Particles” was then used to define ROIs and the mean intensity for the CHMP4B and the RPA2 channels of each ROI were measured. A band-shaped ROI around MN was used to measure mean intensity of the cytoplasm in both channels as regularization factor. For quantification of DOX wash out experiments, cells were washed twice with 1x PBS after 16 hours of DOX incubation (500 ng/ml) and fresh DMEM-F12 complete media without DOX was added to the cells. The same protocol was used for confocal microscopy, western blot and RT-PCR (Extended Data Fig.1d-f). For quantification of the number CHMP4B foci at each time point, the area around nuclei was selected according to DNA staining, and the number of CHMP4B foci was measured for each cell using the Image J/FIJI “Count Maxima” function. Tolerance values were determined for each experiment, depending on the imaging parameters. Three independent experiments were performed.

All data points were plotted using Graphpad Prism.

### Data availability

The datasets generated and/or analyzed during the current study are available from the corresponding authors on reasonable requests.

---

**ORIGINAL WESTERNBLOTS 1.**
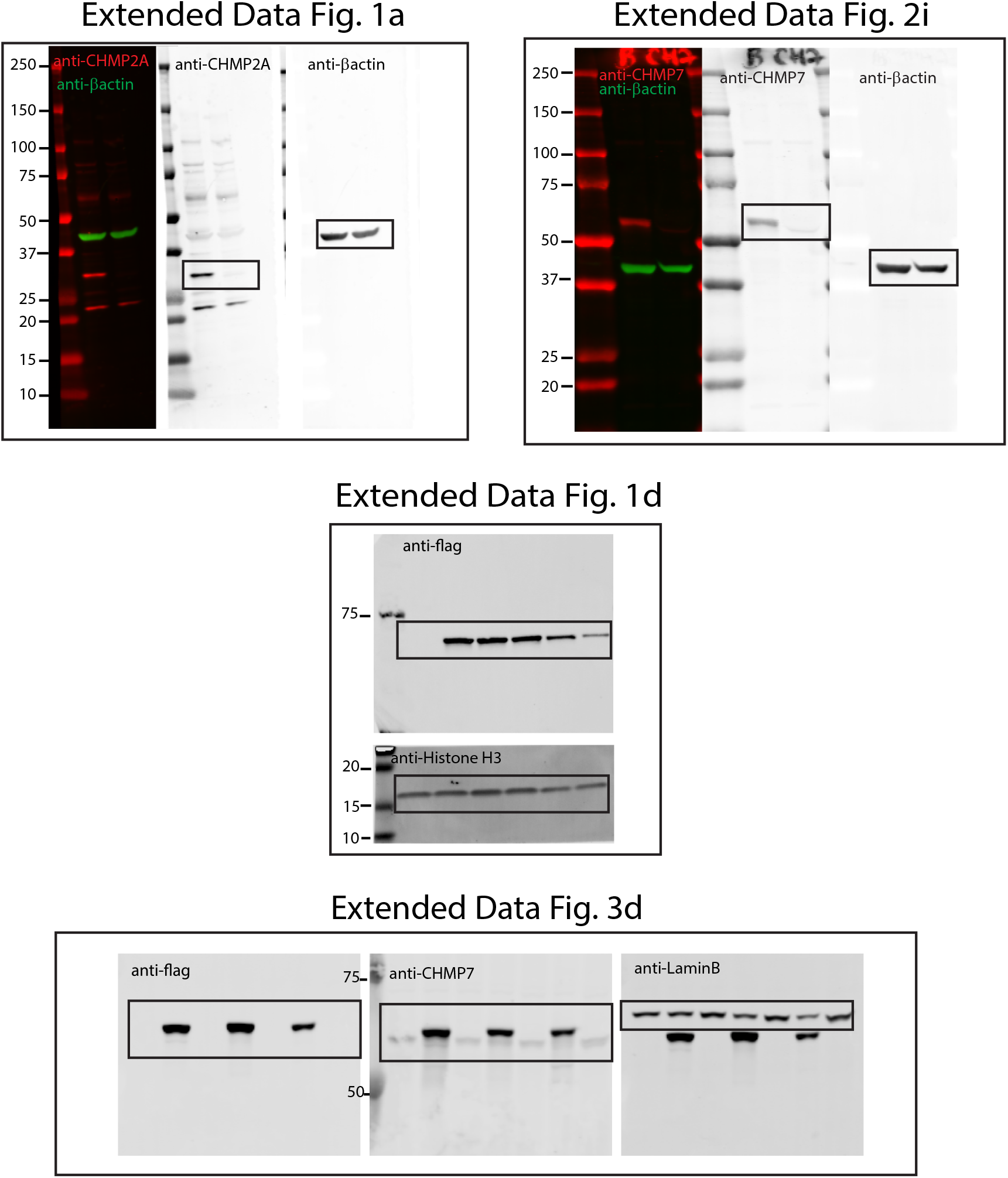

**ORIGINAL WESTERNBLOTS 2.**
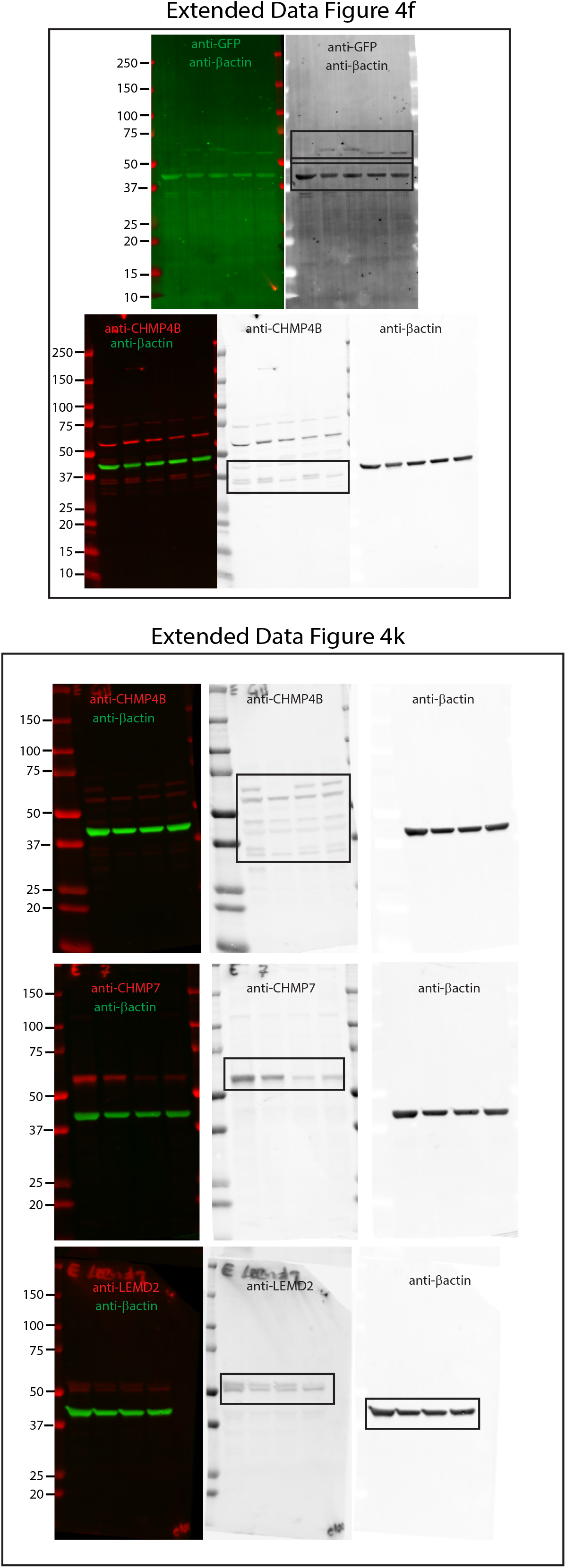

